# Analysis of stress-induced surfaceome remodeling reveals surface accumulation of the cation-independent mannose-6-phosphate receptor (CI-M6PR)

**DOI:** 10.64898/2026.02.26.708128

**Authors:** Fiorella R. Mazzone, Gina Gräßle, Zuzana Storchová, Markus Räschle, Tanja Maritzen

## Abstract

To ensure their survival in the face of stressors, cells have evolved stress response programs. While several transcriptional stress responses have been elucidated, little is known about the impact of stressors on membrane transport and the protein composition of the cell surface. Yet, the dynamic remodeling of the surfaceome by processes such as endocytosis is likely central for the adaptation to stress as it shapes cellular responses by influencing ion uptake and numerous signaling cascades. Indeed, we show that different stressors decrease endocytosis, thereby facilitating cellular adaptation. Using quantitative mass spectrometry, we delineate stress-specific surfaceome alterations in response to osmotic, oxidative and heat stress. Among other adaptive changes, we uncover that osmotic stress leads to a striking surface accumulation of the cation-independent mannose-6-phosphate receptor (CI-M6PR). Mechanistically, we demonstrate that osmotic stress decreases the endocytosis of CI-M6PR while upregulating its lysosomal exocytosis. These results suggest that CI-M6PR might play an important role in the cellular resilience against osmotic stress.

## INTRODUCTION

Cells are frequently faced with alterations in their environment, including changes in temperature, redox state or the osmolarity of the extracellular medium. To ensure their survival, mammalian cells have evolved stress response programs, enabling them to adapt to adverse conditions. System-wide analyses by transcriptomics and proteomics have already provided detailed insights into some aspects of these stress responses such as the transcriptional upregulation of chaperone proteins to assist with protein folding in heat-stressed cells (Alagar Boopathy et al., 2022; Richter et al., 2010). However, the acute effects of different stressors on membrane transport processes and the protein composition of the plasma membrane, i.e. the surfaceome, are much less understood, even though the plasma membrane is likely the first cellular site encountering certain stress conditions. In addition, with its vast complement of transmembrane proteins that range from transporters to nutrient and signaling receptors, the plasma membrane is ideally positioned to preserve ion homeostasis and nutrient uptake and to adapt crucial signaling cascades in response to extracellular cues. Besides, the composition of the plasma membrane itself can be quickly adapted by rerouting proteins from internal membranes to the surface by exocytic processes or by altering the removal of surface proteins by endocytic internalization.

The predominant endocytic mode in many cells is clathrin-mediated endocytosis (CME), where clathrin together with additional endocytic factors causes the invagination of a plasma membrane patch into a 100 nm-sized vesicle that delivers its cargo to the endosomal system (Kaksonen and Roux, 2018). This pathway was proposed to account for ∼95% of total protein endocytic flux (Bitsikas et al., 2014) and, thus, is a prime candidate for modulating the surfaceome upon stress (Lopez-Hernandez et al., 2020a). CME is especially well suited to this task by allowing the selective uptake of specific transmembrane proteins via the recognition of distinct sorting motifs by endocytic adaptor proteins (Azarnia Tehran et al., 2019; Lopez-Hernandez et al., 2020a). In addition, CME occurs on the time scale of minutes and could therefore provide a first line of defense until the slower transcriptional responses take effect. Besides, it might also contribute to the induction of such programs.

In line with this, we have previously shown that a decrease in the endocytosis of the Na^+^/H^+^ exchanger NHE7 (encoded by SLC9A7) upon hyperosmotic stress led to its enrichment at the plasma membrane which plays a crucial role in the resilience of murine astrocytes to osmotic stress by triggering a TFEB/TFE3-dependent increase in the cellular degradative capacity via lysosome biogenesis (Lopez-Hernandez et al., 2020b). This allows osmotically stressed cells to more efficiently cope with the increased load of aggregated proteins which accumulate due to the osmotic insult. Consistently, in absence of NHE7 osmotic stress leads to a greater protein aggregation and a further decrease in cell survival. This example illustrates that endocytosis can play a key role in the intracellular stress response. However, beyond such examples little is known about the extent to which different stress conditions impact the overall endocytic capacity, which changes in the endocytic sorting of individual proteins are induced by stress and how those changes might contribute to a successful cellular adaptation to stress (Lopez-Hernandez et al., 2020a). Previous studies often only measured stress-induced changes in the uptake of one individual endocytic cargo (Lopez-Hernandez et al., 2020a), which does not necessarily reflect stress-induced alterations in bulk endocytic flux and cannot reveal the potentially broad landscape of surfaceome changes. To systematically address the impact of different stressors on endocytosis and the surfaceome, we analyzed on the one hand bulk endocytic uptake upon osmotic, oxidative and heat stress. On the other hand, we employed surface biotinylation of differently stressed cells in combination with quantitative mass spectrometry (MS) to map the surfaceome changes induced by specific stressors. This revealed a general decrease in endocytosis upon stress, uncovered stress-specific surfaceome alterations and identified the cation-independent mannose-6-phosphate receptor (CI-M6PR) as a highly stress-responsive protein which is strongly enriched at the plasma membrane upon osmotic stress, not only due to a decrease in its endocytosis, but also because of increased lysosomal exocytosis.

## RESULTS

### Establishing suitable stress conditions

For our comparative analysis of stress-specific alterations in endocytosis and surfaceome composition, we selected three physiologically relevant stressors at a comparably mild dosage that leads within hours to detectable effects on cellular processes without causing severely increased cell death, thereby leaving a window for stress-induced adaptation. These stressors were initially applied for 1 h to be able to track effects of altered membrane transport without the slower effects of transcriptional regulation as confounding factor.

The first stress condition we chose was hyperosmotic stress since our previous study had already uncovered adaptive changes triggered by this stressor in murine astrocytes (Lopez-Hernandez et al., 2020b). To validate our findings in an independent cellular model, we chose for this study telomerase-immortalized retinal pigment epithelium cells (hTERT-RPE-1) which encounter hyperosmotic stress for example during diabetic retinopathy (Willermain et al., 2014). To establish the right dosage of osmotic stress, we tested 600, 750 and 900 mOsm/kg by adjusting the medium’s osmolarity with D-mannitol, a reagent also used clinically to adjust osmolarity without side effects (Better et al., 1997; Paczynski, 1997; Wijdicks, 2023). As readout we monitored protein aggregation immediately after the 1 h stress period and 24 h later via the dye Proteostat which turns fluorescent upon intercalating into misfolded proteins. While 600 mOsm/kg did not even after 24 h show a significant effect, 750 mOsm/kg led to a significant increase in protein aggregates after 24 h. Since this effect remained comparable at 900 mOsm/kg, but was accompanied by pronounced cell shrinkage and decreased cell viability, we chose 750 mOsm/kg as stress condition (Fig. 1 A,B).

**Fig. 1.**
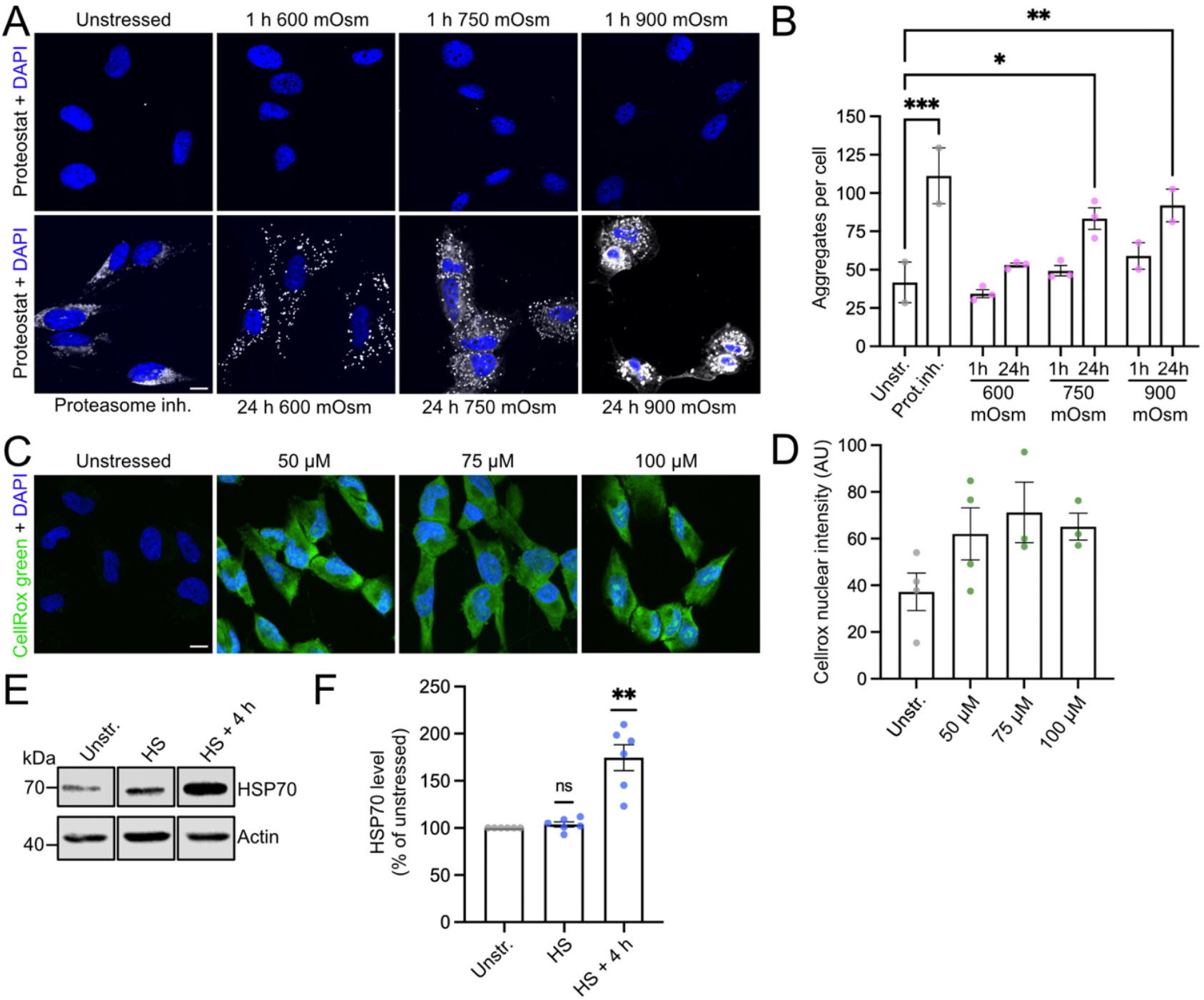
Establishing stress conditions. *A and B*, hTERT-RPE-1 cells were treated at 37°C as indicated, and protein aggregates were visualized by Proteostat (depicted in white). Nuclei were labelled with DAPI (depicted in blue). Proteasome inhibition by 18 h application of 10 µM MG-132 was used as positive control. *A*, representative confocal images. Scale bar, 10 µM. *B*, quantification of the number of protein aggregates. 750 mOsm was the mildest osmotic stress condition to cause a significant increase in protein aggregation after 24 h. Analysis by one-way ANOVA plus Dunnett’s post-test. *C and D*, cells were treated with the indicated concentrations of the oxidative stressor tert-butyl hydroperoxide (TBH) for 1 h at 37°C. Oxidative stress was detected via CellRox fluorescence (depicted in green). Nuclei were labelled with DAPI (depicted in blue). *C*, representative confocal images. Scale bar, 10 µM. *D*, quantification of CellROX green fluorescence. Combined analysis of samples treated with either 50 or 75 µM TBH vs control by unpaired t-test indicates a significant increase in oxidative stress upon TBH treatment (p<0.05). *E and F*, cells were incubated for 1 h at 42°C and either immediately processed for Western blotting (HS) or 4 h later (HS+4h). The induction of the heat stress response was measured by probing for HSP70. Actin was used as loading control. *E*, representative Western blot. Bands originate from same blot. *F*, quantification of HSP70 levels normalized to actin levels and expressed as % of the unstressed control of the respective experiment. Analysis by one sample t-test. *B,D,F*: Columns depict mean ± SEM. Puncta indicate independent experiments. Ns, not significant; *=p<0.05, **=p<0.01, ***=p<0.001. Details on n and p-values are provided in Source Data file.

As second stressor we chose oxidative stress since this is a condition encountered in many human diseases (Barnham et al., 2004; Ebding et al., 2025; Patel, 2016) and was previously reported to influence endocytic trafficking (Cavalli et al., 2001) even though no physiological relevance of this response has been demonstrated so far. We selected tert-butyl hydroperoxide (TBH) to reliably induce oxidative stress, and tested 50, 75 and 100 µM using the fluorescence of CellROX green, a membrane-permeable sensor that emits a strong fluorescent signal upon oxidation, as a readout. After 1 h of treatment, all conditions led to a visible increase in fluorescence (Fig. 1 C,D). Since the effect seemed to saturate at 75 µM, we chose this dosage for our experiments.

The third stressor we selected was heat stress, choosing 42°C as a temperature which is comparable to febrile conditions observed in severe illness and was previously reported to induce a heat shock response within 1 h (Vega et al., 2010; Zhou et al., 2015). We confirmed this in our cellular system by monitoring the known transcriptional upregulation of HSP70 (Zhou et al., 2015) immediately after the 1 h heat stress period and 4 h later.

As expected, immediately after the heat stress the transcriptional response was not yet measurable, however, 4 h later there was a significant increase in HSP70, validating the effectiveness of the treatment (Fig. 1 E,F).

To ensure that all chosen conditions, even upon longer treatment, had only mild effects, we assessed their impact on cell death and proliferation as sensitive readouts for cellular fitness. To test for necrotic cell death, we performed a staining with the membrane-impermeant dye SYTOX after applying the stressors for 4 h or 8 h, which labels cells with compromised membrane integrity. Our results indeed showed that the number of necrotic cells was low in all conditions. Only upon osmotic stress did we observe a significantly higher number of necrotic cells which, however, did not exceed ∼2% of all cells (Fig. 2 A,B). Similarly, the number of apoptotic cells, measured by detecting caspase activity, while being significantly increased after 8 h by all stressors, remained below 2% (Fig. 2 C,D). However, proliferation proved to be highly vulnerable to stress irrespective of the stressor since all stress conditions suppressed cell proliferation while being applied for 24 h (Fig. 2E), making this a sensitive readout for stress effects.

**Fig. 2.**
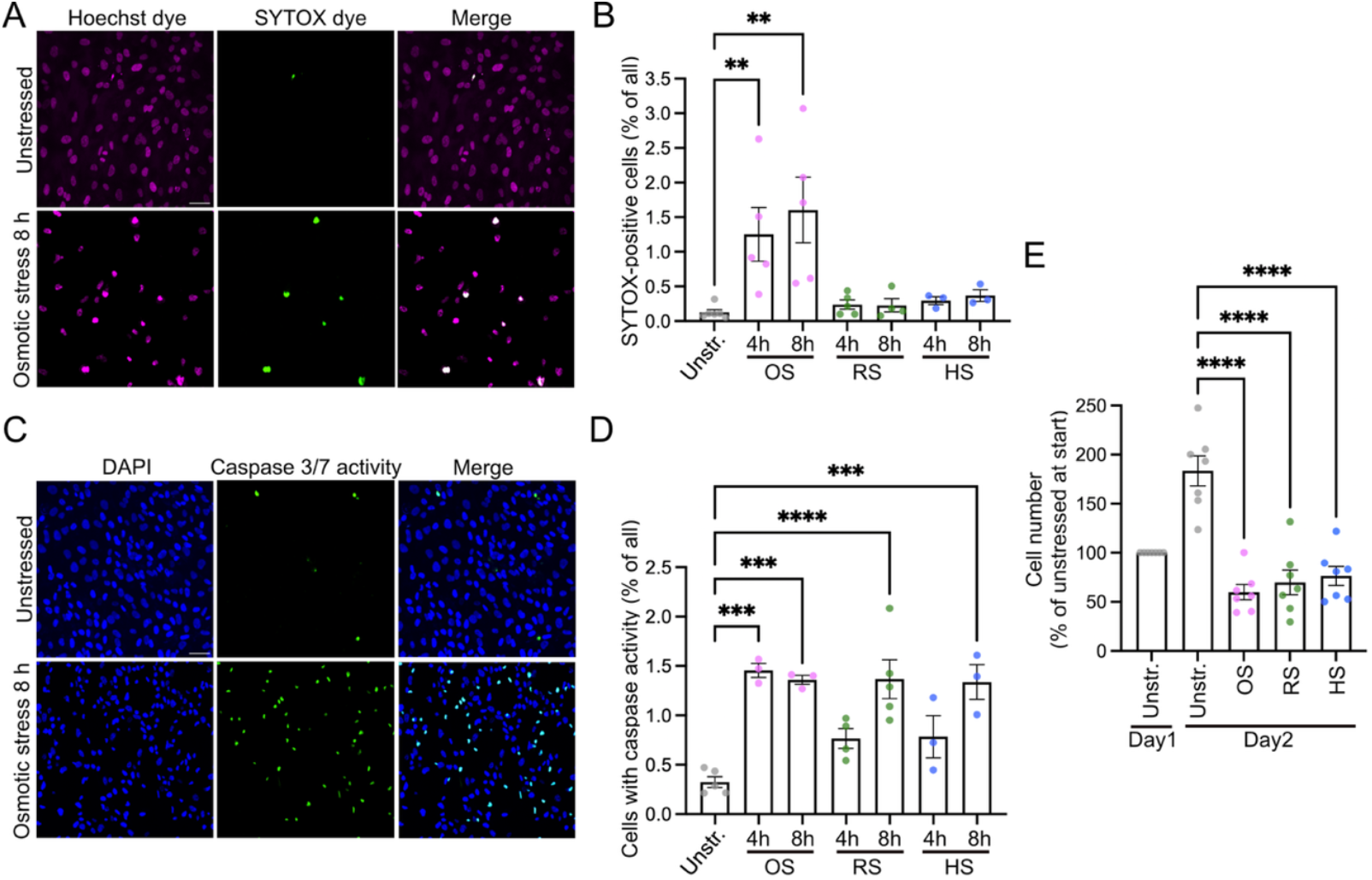
Long-term effects of chosen stress conditions on cell viability and proliferation. *A and B*, cells were left unstressed or subjected for 4 h or 8 h to either osmotic (750 mOsm D-mannitol; OS), oxidative (75 µM TBH; RS) or heat stress (42°C, HS) and subsequently incubated with SYTOX dye to label necrotic cells. Nuclei were labelled with Hoechst dye. *A*, representative confocal images of unstressed cells and cells subjected for 8 h to osmotic stress. Scale bar, 10 µm. *B*, quantification of the percentage of SYTOX-positive nuclei per image for all indicated conditions. Only osmotic stress lead to a small, but significant incrase in necrotic cells. Analysis by Kruskal-Wallis test plus Dunn’s post-test. *C and D*, cells were left unstressed or subjected for 4 h or 8 h to the same stressors as in (A) and subsequently probed for caspase 3/7 activity to label apoptotic cells. Nuclei were labelled with DAPI. *C*, representative confocal images of unstressed cells and cells subjected for 8 h to osmotic stress. Scale bar, 10 µm. *D*, quantification of the percentage of cells exhibiting caspase 3/7 activity per image for all indicated conditions. At 8 h all stressors lead to a small, but significant increase in apoptotic cells. Analysis by one-way ANOVA plus Dunnett’s post-test. *E*, 25 h after seeding (day 1), one well of unstressed cells was trypsinized and counted as point of reference. The other cells were either left untreated or subjected for 24 h to the same stressors as in (A-D) before trypsinization and cell counting on day 2. Cell numbers from day 2 were normalized to the number of unstressed cells of the respective experiment on day 1. While the number of unstressed cells nearly doubled from day 1 to day 2, stressed cells did not increase their numbers. The cell numbers with/without stress on day 2 were analyzed by one-way ANOVA with Dunnett’s post-test. *B, D, E*: Columns depict mean ± SEM. Puncta indicate independent experiments. *=p<0.05, **=p<0.01, ***=p<0.001, ****=p<0.0001. Details on n and p-values are provided in Source Data file.

### Decreased endocytosis upon stress

Having established suitable stress conditions, we went on to evaluate their impact on endocytosis by measuring on the one hand the uptake of fluorescently-labelled transferrin as signature cargo for clathrin-mediated endocytosis. On the other hand, we also determined the bulk uptake of surface proteins since the transferrin receptor itself might be subject to a unique stress-induced endocytic regulation and therefore could be a biased readout. For measuring bulk endocytic internalization, we labeled surface proteins indiscriminately with sulfo-NHS-SS-biotin and allowed their uptake. After cleaving off the remaining surface-bound biotin, we permeabilized the cells and detected the internalized biotinylated proteins by fluorescently-labelled streptavidin.

In all cases, the stressors were applied for 1 h before measuring internalization in a 15 min time window. In line with the mild overall effects of the chosen stressors, none of them abrogated endocytosis (Fig. 3A-D). However, all of them caused a significant decrease in transferrin as well as bulk endocytosis. For transferrin endocytosis, this decrease was in the range of 15-20% for osmotic and heat stress, and about 25% for oxidative stress (Fig. 3 A,B). Likewise, in terms of bulk endocytosis, oxidative stress had the largest impact with a reduction of ∼35%, in contrast to the 10-15% decrease observed for osmotic and heat stress (Fig. 3 C,D). Exemplarily, for osmotic stress we also analyzed the level of EEA1 as marker for early endosomes to which endocytic cargo is delivered. In line with the decrease in endocytosis, the EEA1 level was reduced by nearly 50% after 1 h of exposure to hyperosmotic medium (Fig. 3 E,F) supporting the notion that endocytic trafficking is downregulated under stress.

**Fig. 3.**
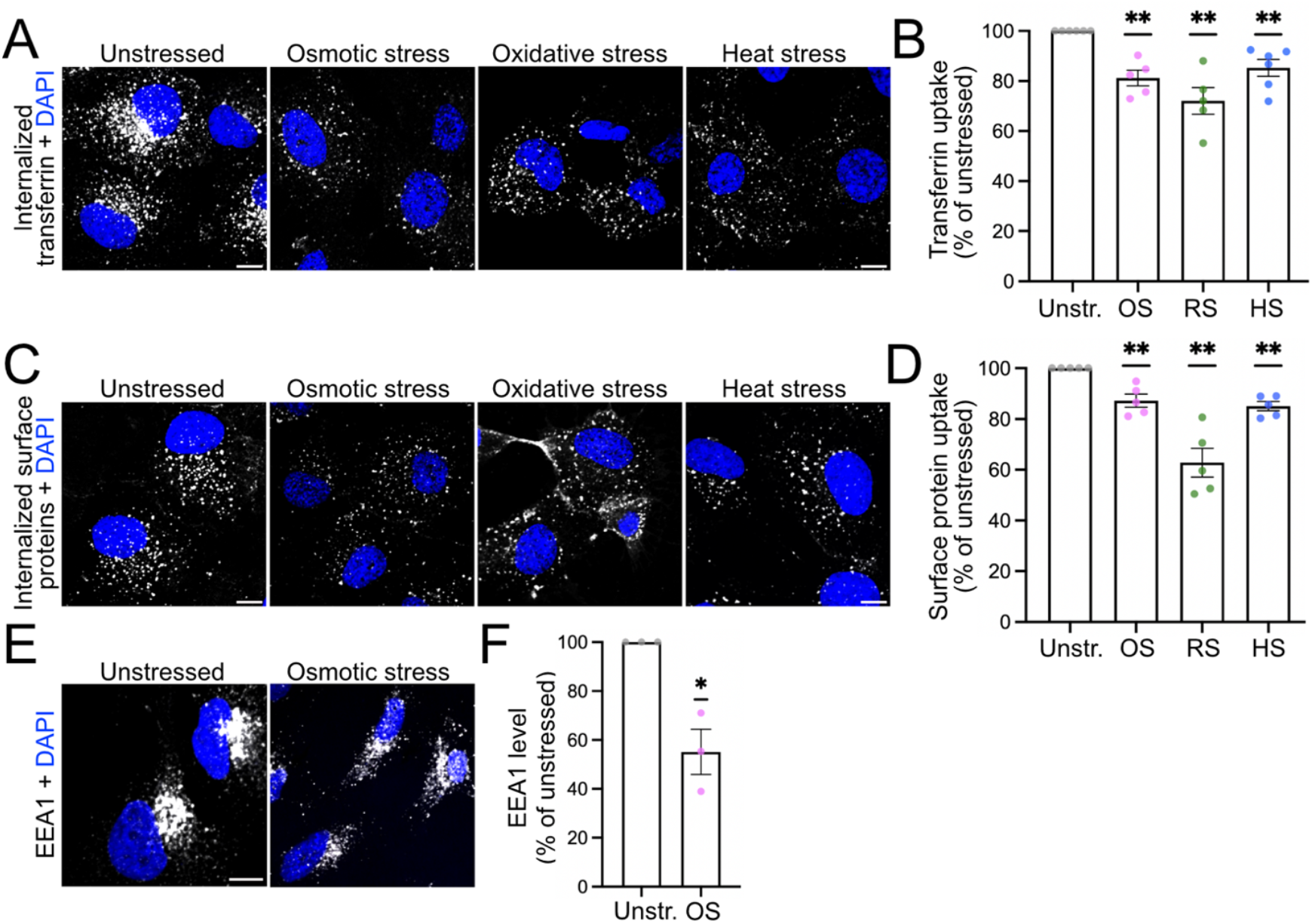
Decreased endocytosis upon stress. *A-D*, cells were either left unstressed or subjected for 1 h to either osmotic (750 mOsm D-mannitol; OS), oxidative (75 µM TBH; RS) or heat stress (42°C, HS). Immediately afterwards, cell surface proteins were biotinylated, and cells then shifted in the presence of transferrin to 37°C for 15 min to allow for endocytosis. Subsequently, internalized proteins were detected via fluorescently-labeled streptavidin, and nuclei were labelled with DAPI (depicted in blue). *A and C*, representative confocal images of internalized transferrin (A) or internalized surface proteins (C). Scale bar, 10 µm. *B and D*, analysis of transferrin uptake (B) and surface protein uptake (D) via quantification of fluorescent puncta per cell normalized to the unstressed condition of the respective experiment. Statistical evaluation via one sample t-tests shows that all stress conditions reduce endocytic uptake. *E and F*, cells were either left unstressed or subjected for 1 h to osmotic stress (750 mOsm D-mannitol; OS). Fixed cells were stained with EEA1-specific antibodies. Nuclei were labeled with DAPI (depicted in blue). Scale bar, 10 µm. *E*, representative confocal images. *F*, quantification of EEA1 fluorescence intensity per cell normalized to the unstressed condition of the respective experiment. Statistical evaluation via one sample t-test reveals lower EEA1 levels upon osmotic stress. *B, D, F*: Columns depict mean ± SEM. Puncta indicate independent experiments. *=p<0.05, **=p<0.01. Details on n and p-values are provided in Source Data file.

### Stress-specific changes in the composition of the surfaceome

Even though endocytic internalization in general is affected by stress, it is likely that the uptake of individual surface proteins with their unique endocytic regulation might be affected to different extents by different stress conditions. To probe this hypothesis in an unbiased manner, we performed surface biotinylations of unstressed and stressed cells followed by a quantitative MS-based analysis of the isolated surfaceome (Supplementary Tables 1 and 2). The principal component analysis and heatmaps of our data demonstrated consistency between replicates and a clear separation of our controls (unstressed cells plus/minus addition of biotin) from the stress conditions and also between the osmotic stress and heat stress condition (Supplementary Fig. 1 A-D; the oxidative stress condition could not be directly compared since it was analyzed in an independent MS run). This indicated already that the different stress conditions lead to changes in the surfaceome that are unique to the specific stressor. This is also reflected in the volcano plots depicting the significant changes in the surface abundance of individual proteins observed upon the three different stress conditions (Fig. 4 A-C). To rule out that the quantified surfaceome alterations are an indirect consequence of altered protein abundance, we also analyzed the whole-cell proteome of unstressed and stressed cells by quantitative MS (Supplementary Table 3). Principal component analysis again showed a clear separation between the proteomes of unstressed and stressed cells (Supplementary Figure 2A). In addition, the results confirmed that the candidate proteins with increased surface abundance which we followed up on (as far as they had been detected in the whole cell proteome analysis), were not just enriched at the surface because of a significantly increased overall abundance (Supplementary Figure 2 D-F). The only exception was the SPPL2A expression upon heat stress which appeared somewhat increased. However, this increase was not confirmed by subsequent immunoblotting (Figure 5 I). In addition, we checked the whole-cell proteome dataset for the expression levels of 28 canonical endocytic proteins to investigate whether any of these might show consistently decreased levels as potential mechanistic reason for the observed decrease in endocytosis. However, we did not detect any consistent changes across the different stress conditions (Supplementary Figure 2 G-I and Supplementary Table 4). This is consistent with the more likely hypothesis that the endocytic uptake of individual proteins is rather altered due to post-translational modifications in endocytic or cargo proteins which affect their complex formation. Interestingly, we detected 37 proteins whose levels were downregulated and 14 proteins whose levels were upregulated upon all three stressors (Supplementary Figure 2 B,C and Supplementary Table 5) suggesting that there might be cellular processes that are subject to a coordinated regulation across different stress conditions.

**Fig. 4.**
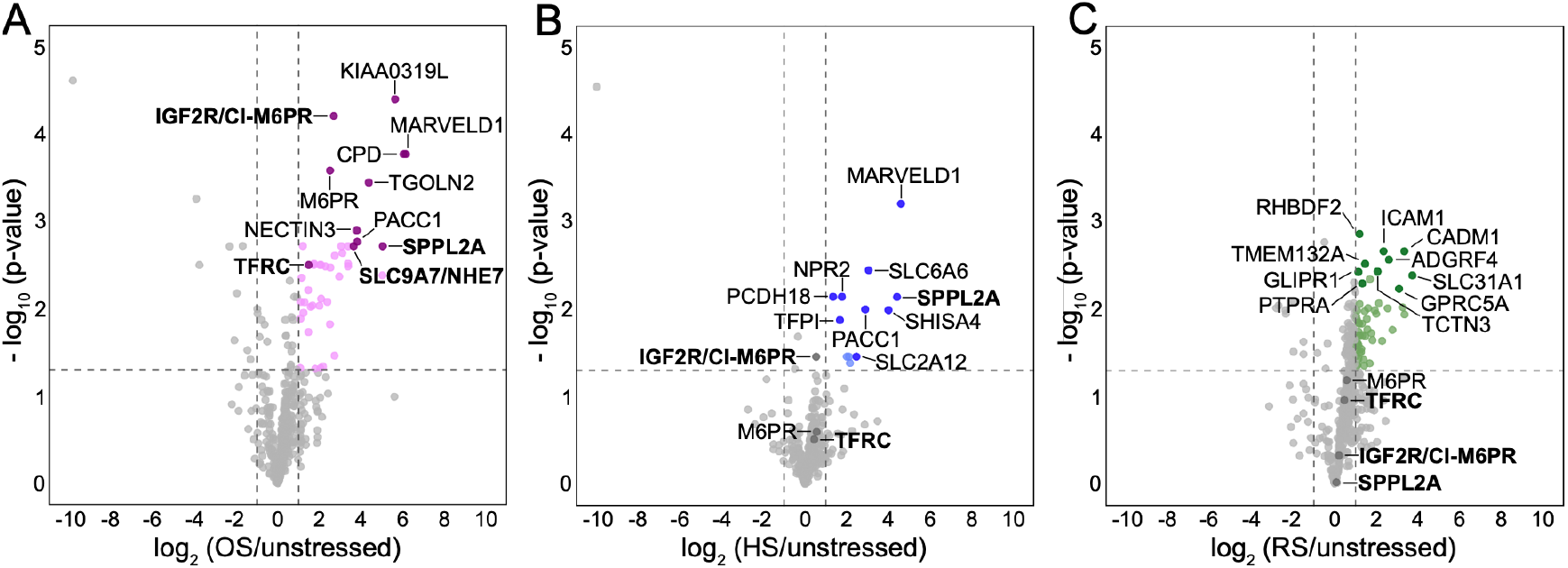
Stress-induced surfaceome alterations. *A*-*C*, volcano plots of log_2_-fold changes and −log_10_-transformed, Benjamini– Hochberg corrected p-values, representing surfaceome alterations induced by osmotic stress (A), heat stress (B) or oxidative stress (C), with significantly surface-enriched candidates represented by coloured puncta and validated proteins in bold (see also Supplementary Tables 1 and 2). Dashed lines represent the thresholds for statistically significant changes in protein abundance (threshold BH-adjusted p-value<0.05, |log_2_FC|>1).

**Fig. 5.**
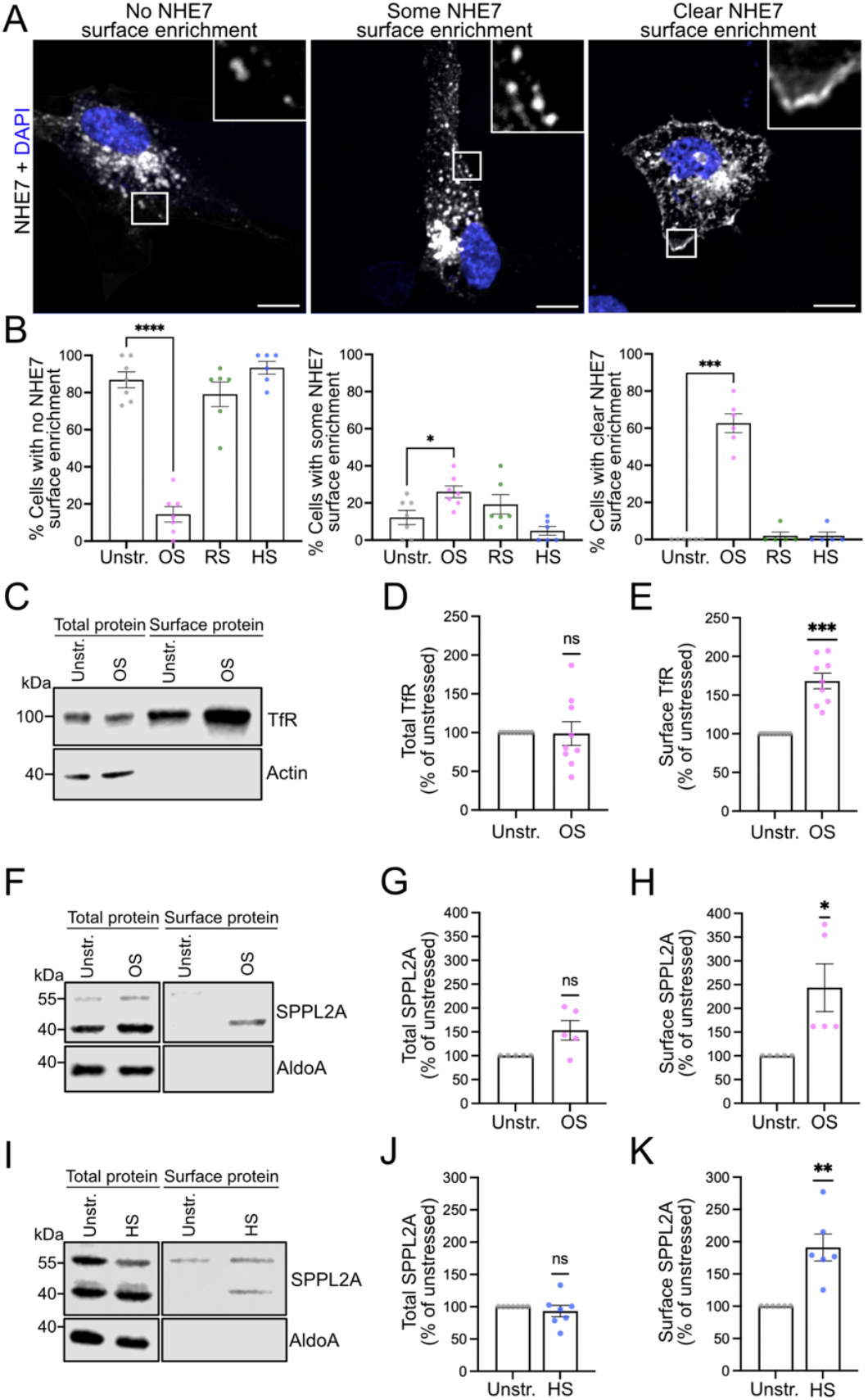
Validation of stress-induced surfaceome alterations. *A and B*, NHE7 surface accumulation upon osmotic stress. Cells transfected with NHE7-mCherry were left unstressed or subjected for 1 h to either osmotic (750 mOsm D-mannitol; OS), oxidative (75 µM TBH; RS) or heat stress (42°C, HS). Nuclei of fixed cells were labelled with DAPI (depicted in blue). *A*, representative confocal images showing cells with different degrees of NHE7 surface enrichment matching the categories used for quantification in (B). Scale bar, 10 µm. *B*, quantification of the percentage of cells showing either no, some or clear surface enrichment upon the different stress conditions. Analysis by one-way ANOVA plus Dunnett’s post-test (left and middle graph) or Kruskall-Wallis test plus Dunn’s post-test (right graph) reveals significant increase in cells with clear NHE7 surface enrichment specifically upon osmotic stress. *C-E*, transferrin receptor surface accumulation upon osmotic stress. Cells unstressed or exposed to 1 h of osmotic stress (750 mOsm D-mannitol; OS) were subjected to surface biotinylation. Total protein samples and surface protein eluates were analyzed by Western blotting and probed for transferrin receptor and aldolase A as control for the selective biotinylation of surface proteins. *C*, representative Western blot. *D*, quantification of total transferrin receptor levels normalized to aldolase A levels, expressed as % of unstressed condition and analyzed by one sample t-test. *E*, quantification of surface transferrin receptor levels expressed as % of unstressed condition. Analysis by one-sample t-test reveals significant increase in surface transferrin receptor level upon osmotic stress. *F-K*, SPPL2A surface accumulation upon osmotic (F-H) and heat stress (I-K). Cells unstressed or exposed to 1 h of osmotic stress (F-H; 750 mOsm D-mannitol; OS) or heat stress (I-K, 42°C, HS) were subjected to surface biotinylation. Total protein samples and surface protein eluates were analyzed by Western blotting and probed for SPPL2A and aldolase A. *F and I*, representative Western blots. Bands originate from same blots. *G and J*, quantification of total SPPL2A levels normalized to aldolase A levels, expressed as % of unstressed condition and analyzed by one sample t-test. *H and K*, quantification of surface SPPL2A levels expressed as % of unstressed condition. Analysis by one-sample t-test reveals significant increase in surface SPPL2A level upon osmotic and heat stress. *B, D, E, G, H, J, K*: Columns depict mean ± SEM. Puncta indicate independent experiments. Ns, not significant; *=p<0.05, **=p<0.01, ***=p<0.001, ****=p<0.0001. Details on n and p-values are provided in Source Data file.

### Confirmation of NHE7 as stress-responsive protein

We started our validation of individual proteins which accumulate at the plasma membrane in a stress-dependent manner by searching the surfaceome datasets for NHE7 which we had previously found enriched at the surface of osmotically stressed murine astrocytes (Lopez-Hernandez et al., 2020b). In support of the reliability of our previous findings and the quality of our new surfaceome data, NHE7 was identified as surface-enriched upon osmotic stress (Fig. 4A). To confirm this finding for hTERT-RPE-1 cells by an independent experimental approach, we subjected cells expressing a fluorescently-tagged version of NHE7 (in absence of a reliable antibody) to the different stress conditions and scored in a blinded manner whether the transfected cells displayed NHE7 at the membrane (Fig. 5 A,B). This analysis indicated not only that osmotic stress leads to a highly significant increase in the number of cells with clear NHE7 surface enrichment, but also revealed that this effect is highly specific for osmotic stress underlining that the different stress conditions can affect individual proteins in a highly selective manner.

### Validation of additional candidates including CI-M6PR

For the further validation, we concentrated on the osmotic and heat stress condition since there was some overlap between the two in regards to the specific surface-enriched proteins as detailed below. A second protein which our proteomics data revealed to be selectively accumulating at the surface upon osmotic stress and which we successfully validated was the transferrin receptor (Fig. 4A, Fig. 5 C-E). These data are consistent with our earlier observation of decreased transferrin endocytosis upon osmotic stress (Fig. 3B) since an impairment of transferrin receptor internalization will at the same time lead to its surface accumulation and a decrease in the uptake of transferrin. In addition, we confirmed that the protein signal peptide peptidase-like 2A (SPPL2A; Fig. 4A) which is normally present in late endosomes and lysosomes (Behnke et al., 2011) was also enriched at the plasma membrane upon osmotic stress (Fig. 5 F-G). Interestingly, SPPL2A accumulated also verifiably at the surface upon heat stress (Fig. 4B, Fig. 5I-K). Finally, another protein reproducibly accumulating at the membrane upon osmotic and heat stress was the cation-independent mannose-6-phosphate receptor (CI-M6PR), also known as insulin-like growth factor 2 receptor (IGF2R) (Fig. 4 A,B; Fig. 6 A-F). This receptor caught our interest due to its striking surface-enrichment upon osmotic stress (Fig 4A, Fig. 6F) which motivated us to further elucidate the characteristics of its membrane enrichment and the underlying mechanism.

**Fig. 6.**
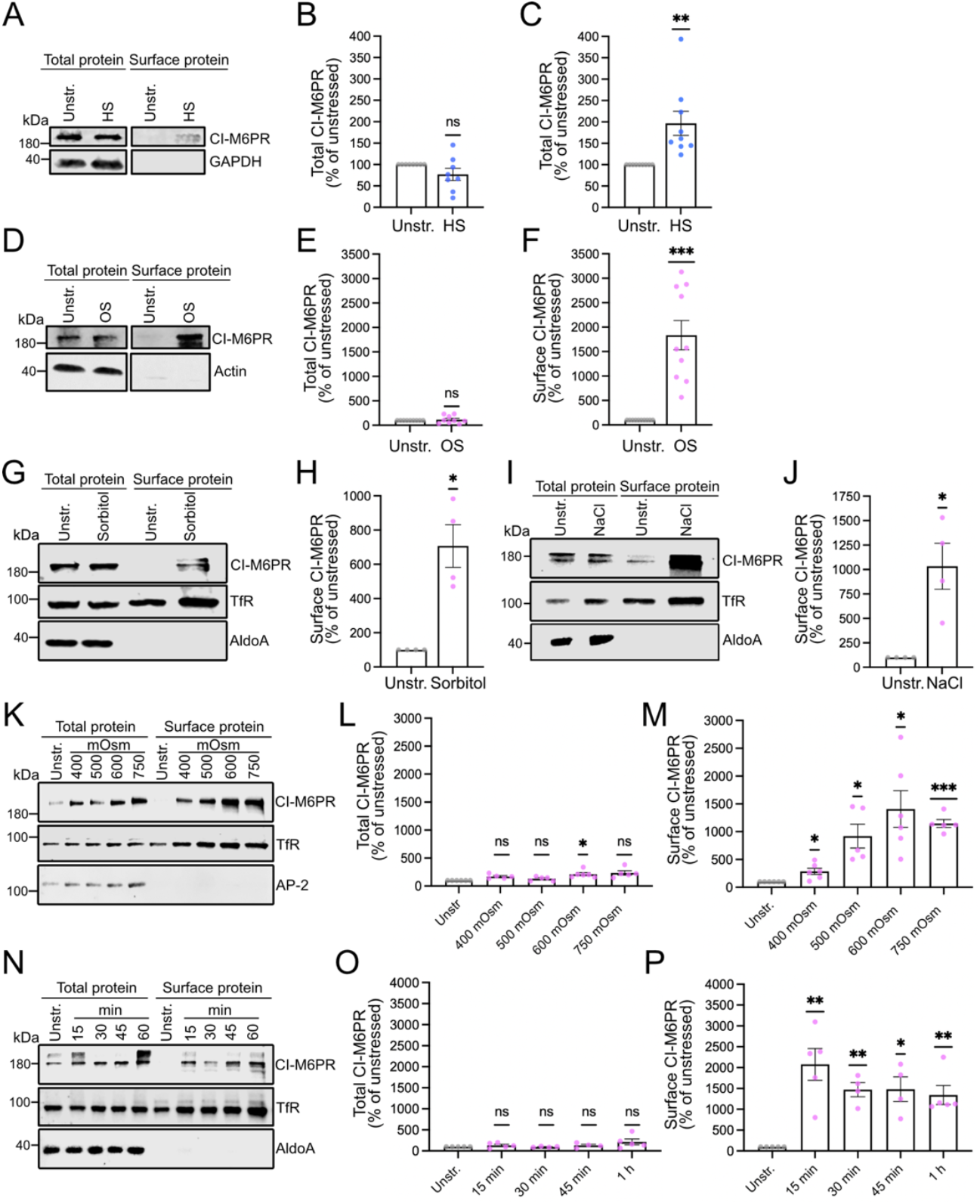
Reliable and fast accumulation of CI-M6PR upon osmotic stress. *A-F*, CI-M6PR surface accumulation upon heat (A-C) or osmotic (D-F) stress. Cells unstressed or exposed for 1 h to heat stress (A-C, 42°C, HS) or osmotic stress (D-F, 750 mOsm D-mannitol; OS) were subjected to surface biotinylation. Total protein samples and surface protein eluates were analyzed by Western blotting and probed for the indicated proteins. *A and D*, representative Western blots. Bands originate from same blots. *B and E*, quantification of total CI-M6PR levels normalized to loading control levels, expressed as % of unstressed condition and analyzed by one sample t-test. *C and F*, quantification of surface CI-M6PR levels expressed as % of unstressed condition. Analysis by Wilcoxon signed rank test (C) or one-sample t-test (F) reveals significant increase in surface CI-M6PR level upon heat and osmotic stress. *G-J*, osmotic stress via sorbitol or NaCl addition also induces CI-M6PR surface accumulation. Cells unstressed or exposed for 1 h to 750 mOsm sorbitol (G,H) or NaCl (I,J) were subjected to surface biotinylation. Total protein samples and surface protein eluates were analyzed by Western blotting and probed for CI-M6PR, transferrin receptor and aldolase A. *G and I*, representative Western blots. *H and I*, quantification of surface CI-M6PR levels expressed as % of unstressed condition. Analysis by one-sample t-test (F) reveals significant increase in surface CI-M6PR level upon sorbitol or NaCl. *K-M*, CI-M6PR surface accumulation already induced with lower osmolarities. Cells unstressed or exposed for 1 h to the indicated hypertonic solutions were subjected to surface biotinylation. Total protein samples and surface protein eluates were analyzed by Western blotting and probed for CI-M6PR, transferrin receptor and AP-2. *K*, representative Western blots. *L*, quantification of total CI-M6PR levels normalized to AP-2 levels, expressed as % of unstressed condition and analyzed by Wilcoxon signed rank test. *M*, quantification of surface CI-M6PR levels expressed as % of unstressed condition. Analysis by one-sample t-test (F) reveals significant increase in surface CI-M6PR level already upon 400 mOsm. *N-P*, CI-M6PR accumulates quickly at surface. Cells unstressed or exposed for the indicated times to 750 mOsm D-mannitol were subjected to surface biotinylation. Total protein samples and surface protein eluates were analyzed by Western blotting and probed for CI-M6PR, transferrin receptor and aldolase A. N, representative Western blots. *O*, quantification of total CI-M6PR levels normalized to aldolase A levels, expressed as % of unstressed condition and analyzed by one sample t-test. *P*, quantification of surface CI-M6PR levels expressed as % of unstressed condition. Analysis by one-sample t-test reveals significant increase in surface CI-M6PR level already within 15 min. *B,C,E,F,H,J,L,M,O,P*: Columns depict mean ± SEM. Puncta indicate independent experiments. Ns, not significant. *=p<0.05, **=p<0.01, ***=p<0.001. Details on n and p-values are provided in Source Data file.

### Characterization of CI-M6PR surface accumulation

The best understood function of CI-M6PR is to transport lysosomal hydrolases from the trans-Golgi network (TGN) to late endosomes/lysosomes (Gauthier et al., 2024). While CI-M6PR normally shuttles between TGN and late endosomes, a small pool of less than 10% is also present at the plasma membrane (Klumperman et al., 1993) where CI-M6PR can retrieve lysosomal hydrolases which have been mistakenly released from the cell after entering the constitutive secretory pathway (Gauthier et al., 2024; Oshima et al., 1988). In addition, it can interact there with one of the many extracellular ligands it has been reported to bind (Gauthier et al., 2024) including IGF2 (Bohnsack et al., 2024; Morgan et al., 1987), heparanase (Wood and Hulett, 2008), granzyme B (Motyka et al., 2000), uPAR (Godár et al., 1999; Schiller et al., 2009), plasminogen (Godár et al., 1999), latent TGF*β* (Godár et al., 1999) and retinoic acid (Kang et al., 1997). However, the function of most of these interactions is not as well understood.

Compared to the low amount of CI-M6PR usually present at the plasma membrane in unstressed cells, which is hardly detectable in immunoblots of surface biotinylations, there is a small, but significant increase upon heat shock and a tremendous increase upon osmotic stress (Fig. 6A-F). To make sure that this increase was not specific to the use of D-mannitol as osmolyte, we also tested the effects of 750 mOsm/kg hyperosmotic buffer using NaCl or sorbitol as alternative osmolytes. In both conditions, we observed a large and significant increase in the CI-M6PR surface level indicating that its surface accumulation is a general response to osmotic stress independent of the osmolyte used (Fig. 6 G-J). Next, we sought to determine the threshold of hyperosmotic stress at which CI-M6PR surface enrichment starts to occur. To assess this, we tested a series of osmolarities with the lowest being 400 mOsm/kg. Interestingly, 400 mOsm/kg was already sufficient to trigger a significant elevation of CI-M6PR at the plasma membrane (Fig. 6 K-M). However, 500 and 600 mOsm/kg substantially increased the extent of the surface accumulation, but beyond that the effect appeared to be saturated (Fig. 6 K-M). Since membrane transport events occur in the range of minutes, we also tested how rapidly the surface accumulation of CI-M6PR occurs, choosing 15 min as the shortest time window. Consistent with being caused by alterations in endo- or exocytosis, the surface enrichment was already evident at 15 min and as pronounced at this time as at later time points (Fig. 6 N-P), making this an early occurring stress response.

### A decrease in the endocytosis of CI-M6PR promotes its stress-dependent surface enrichment

CI-M6PR is known to be endocytosed by clathrin-mediated endocytosis (Willingham et al., 1981). To establish whether a decreased endocytosis of CI-M6PR could potentially be the cause for its surface accumulation, we determined CI-M6PR surface levels after blocking CME with the endocytosis inhibitor Pitstop2 (von Kleist et al., 2011) which efficiently abrogates transferrin uptake (Fig. 7A). As expected, the inhibition of CME lead to an increase of CI-M6PR at the plasma membrane, even though the level reached was lower than upon osmotic stress (Fig. 7 B,C), supporting the notion that the stress-induced redistribution of CI-M6PR to the plasma membrane involves a CME-dependent mechanism. To probe whether the endocytosis of CI-M6PR is indeed impaired upon osmotic stress we performed an antibody uptake experiment which revealed a similar decrease of CI-M6PR uptake by 25-30% upon heat and osmotic stress (Fig. 7 D,E). Since the internalization of CI-M6PR is mediated via the recognition of the endocytic sorting motif YSKV in its cytoplasmic tail by the endocytic adaptor protein AP-2 (Ghosh et al., 2003; Jadot et al., 1992), we reasoned that the intracellular tail should contain all necessary determinants for causing the stress-induced surface accumulation of CI-M6PR. To test this hypothesis, we expressed GFP fused to the transmembrane and intracellular region of CI-M6PR in hTERT-RPE1 cells and evaluated its redistribution upon osmotic stress. In line with the behaviour of full-length CI-M6PR as reported in the literature (Anitei et al., 2014; Simonetti et al., 2019), the chimeric protein localized mostly to the TGN (Fig. 7F) since also sorting signals for its transport to the TGN are contained in the CI-M6PR cytoplasmic tail (Ghosh et al., 2003). For quantifying a potential stress-induced surface accumulation of the chimera, we used a decrease in the colocalization with the TGN as easy readout for its redistribution, employing Pitstop2 as a positive control. Indeed, osmotic stress and Pitstop both induced a loss of the chimera from the TGN (Fig. 7 F,G), presumably due to a lack of endocytic retrieval of surface-stranded chimeric proteins.

**Fig. 7.**
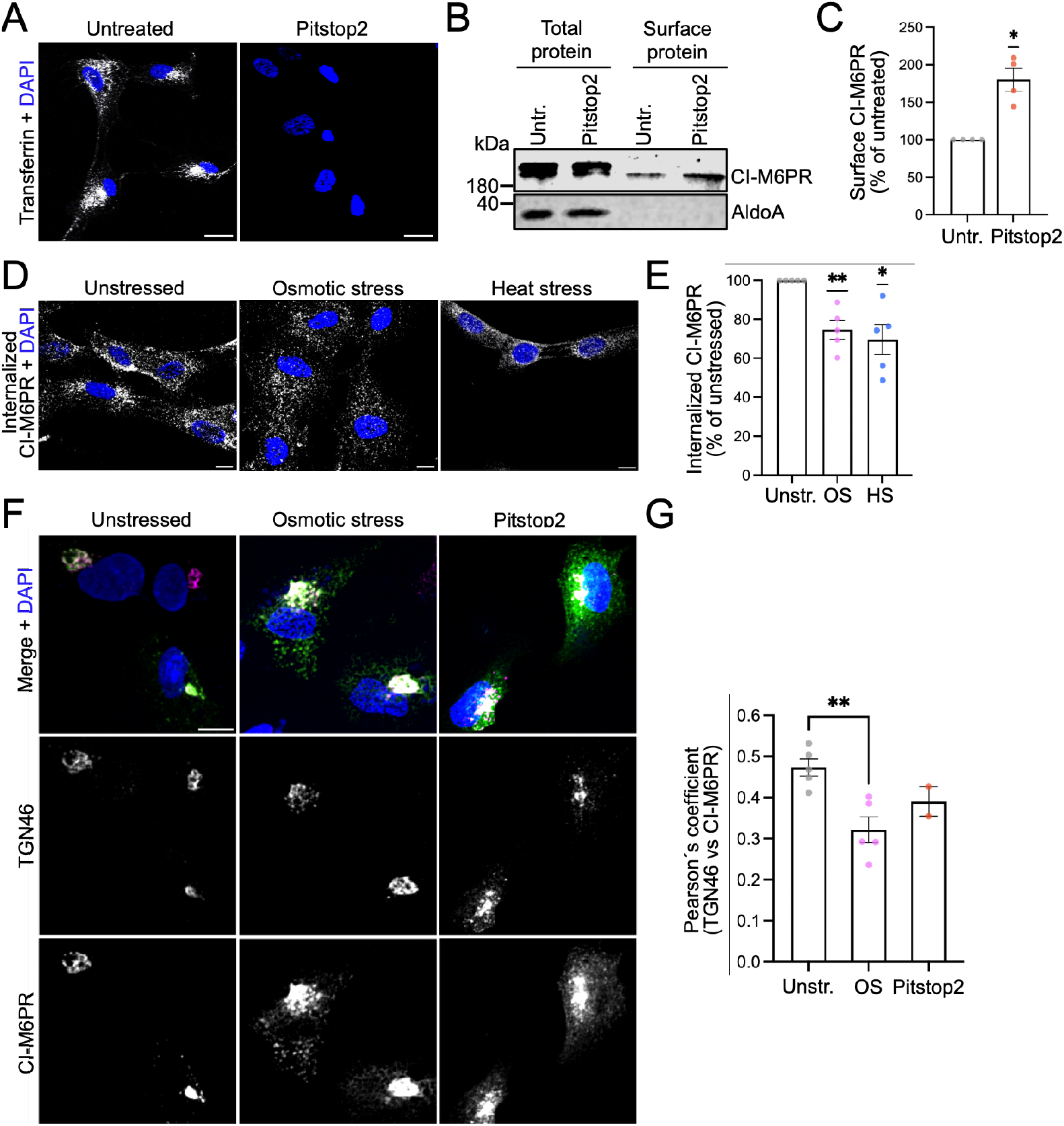
Stress-induced decrease in endocytosis constributes to CI-M6PR surface accumulation. *A-C*, inhibition of clathrin-mediated endocytosis (CME) mimics CI-M6PR surface accumulation upon stress. *A*, cells were incubated for 15 min at 37°C with 30 µM of the endocytosis inhibitor Pitstop2 and 25 µg/ml fluorescently labelled transferrin. Nuclei of fixed cells were visualized by DAPI (in blue). Representative confocal images showing that Pitstop2 is able to block transferrin endocytosis. Scale bar, 10 µm. *B*, cells untreated or exposed to Pitstop2 were subjected to surface biotinylation. Total protein samples and surface protein eluates were analyzed by Western blotting and probed for CI-M6PR and aldolase A. *C*, quantification of surface CI-M6PR levels expressed as % of untreated condition. Analysis by one-sample t-test reveals significant increase in surface CI-M6PR level upon inhibition of CME. *D and E*, CI-M6PR endocytosis is impaired upon stress. *D*, representative confocal images of an antibody feeding assay probing the internalization of CI-M6PR after 1 h of osmotic (750 mOsm D-mannitol) or heat stress (42°C). Nuclei of fixed cells were visualized by DAPI (in blue). Scale bar, 10 µm. *E*, quantification of internalized CI-M6PR expressed as % of unstressed condition. Analysis by one-sample t-test reveals significantly decreased endocytosis upon osmotic and heat stress. *F and G*, intracellular CI-M6PR tail decisive. Cells expressing a EGFP-tagged CI-M6PR variant lacking the extracellular domains were left unstressed or exposed for 1 h to 750 mM D-mannitol or 30 µM Pitstop2. Fixed cells were labeled with TGN46-specific antibodies (in magenta). Nuclei were visualized by DAPI (in blue). Scale bar, 10 μm. *F*, representative confocal images. *G*, quantification of TGN localization of CI-M6PR chimera via measurement of Pearson’s colocalization coefficient. Analysis by one-way ANOVA plus Dunnett’s post-test shows that CI-M6PR lacking its extracellular domains still becomes redistributed upon osmotic stress. *C, E, G*: Columns depict mean ± SEM. Puncta indicate independent experiments. Ns, not significant; *=p<0.05, **=p<0.01. Details on n and p-values are provided in Source Data file.

Therefore, we conclude that a decline in CME, dependent on determinants in the intracellular tail of CI-M6PR, contributes to the osmotic and heat stress-dependent surface accumulation of CI-M6PR.

### Elevated lysosomal exocytosis additionally contributes to the osmotic stress-dependent surface accumulation of CI-M6PR

Even though decreased endocytosis promotes the surface accumulation of CI-M6PR, the comparably small extent of its surface enrichment upon a full inhibition of endocytosis by Pitstop2 was puzzling in light of the much stronger redistribution caused by osmotic stress, arguing for a second mechanism contributing to the accumulation of CI-M6PR at the plasma membrane. Interestingly, when scrutinizing the list of proteins displaying surface enrichment upon osmotic stress, we noticed that there were many late endosomal/lysosomal transmembrane proteins (e.g. cation-dependent (CD)-M6PR, a V-ATPase subunit and LAMP1) among the hits (Fig. 8A). Since not all of them are known to undergo endocytosis, we wondered about an alternative mechanism that might lead to their concerted stress-induced surface enrichment.

**Fig. 8:**
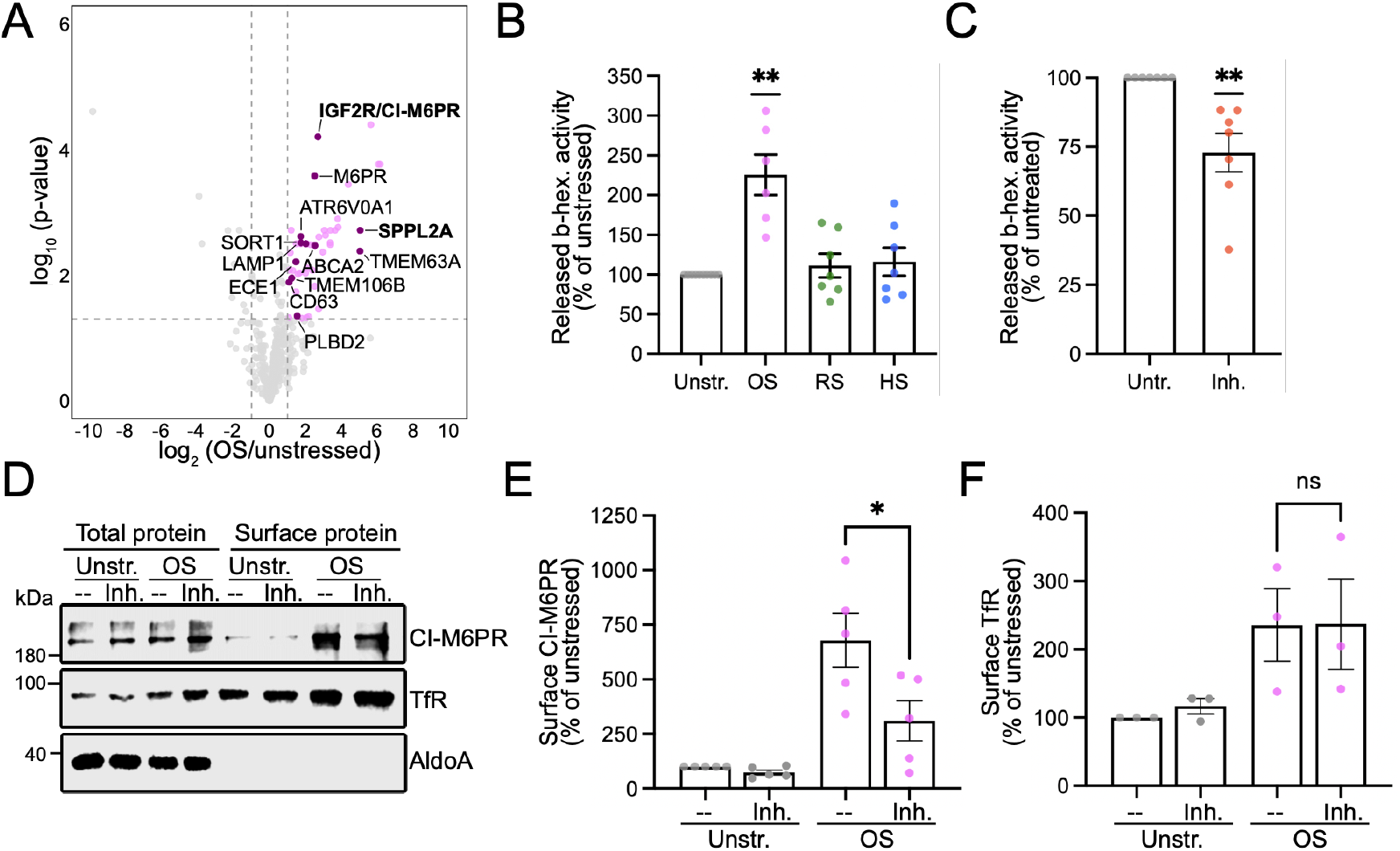
Osmotic stress-induced lysosomal exocytosis contributes to CI-M6PR surface accumulation. *A*, volcano plot of log_2_-fold changes and −log_10_-transformed, Benjamini–Hochberg corrected p-values showing the surface enrichment of lysosomal proteins upon osmotic stress, with significantly surface-enriched candidates in magenta and significantly surface-enriched lysosomal proteins in purple. Dashed lines represent the thresholds for statistically significant changes in protein abundance (threshold BH-adjusted p-value<0.05, |log_2_FC|>1). *B*, osmotic stress induces release of lysosomal enzymes. The activity of the lysosomal enzyme *β*-hexosaminidase was measured in the supernatants of unstressed cells and cells exposed for 1 h to osmotic (750 mOsm D-mannitol; OS), oxidative (75 µM TBH; RS) or heat stress (42°C; HS) and expressed as % of the unstressed condition. Analysis by one-sample t-test shows that specifically OS induces lysosomal exocytosis. *C*, TRPML1 inhibition decreases lysosomal exocytosis. *β*-Hexosaminidase activity was measured in the supernatants of untreated cells and cells treated for 1 h with the TRPML1 inhibitor ML-SI3 and expressed as % of the untreated condition. Analysis by one-sample t-test shows a significant decrease in the release of the lysosomal enzyme upon ML-SI3 treatment. *D-F*, decreased CI-M6PR surface accumulation upon inhibition of lysosomal exocytosis. Cells unstressed or exposed for 1 h to 750 mOsm D-mannitol in presence of 10 µM ML-SI3 or an equal volume of DMSO as vehicle control were subjected to surface biotinylation. Total protein samples and surface protein eluates were analyzed by Western blotting and probed for CI-M6PR, transferrin receptor and aldolase A. *D*, representative Western blots. *E and F*, quantification of surface CI-M6PR (E) and transferrin receptor (F) levels expressed as % of unstressed condition. Comparison of surface accumulation between ML-SI3 treated and untreated osmotically stressed cells by unpaired t-test reveals a significant decrease in the surface accumulation specifically for CI-M6PR upon inhibition of lysosomal exocytosis. *B, C, E, F*: Columns depict mean ± SEM. Puncta indicate independent experiments. Ns, not significant; *=p<0.05, **=p<0.01. Details on n and p-values are provided in Source Data file.

While lysosomal exocytosis is clearly a mechanism expected to lead to a concerted increase of lysosomal proteins at the plasma membrane, we found no indication in the literature of its upregulation upon osmotic stress, while a regulation by oxidative stress had been reported for HeLa cells (Ravi et al., 2016). To clarify the situation for hTERT-RPE-1 cells, we applied the different stressors and measured *β*-hexosaminidase activity in the supernatant as established readout for lysosomal exocytosis (Laulagnier et al., 2011; Rodriguez et al., 1997).

Our results clearly showed that in hTERT-RPE-1 cells osmotic stress was the only stressor which induced lysosomal exocytosis significantly and to a substantial extent (Fig. 8B), suggesting that lysosomal exocytosis could indeed contribute to the surface accumulation of CI-M6PR. To test this hypothesis further, we verified the efficacy of a published inhibitor of lysosomal exocytosis, the TRPML1 antagonist ML-SI3 (Leser et al., 2021), in our cellular model (Fig. 8C). Afterwards, we used this inhibitor to analyze the contribution of lysosomal exocytosis to the surface accumulation of CI-M6PR. As expected, the inhibitor did not have any effect on the anyhow low surface level of CI-M6PR in unstressed cells (Fig. 8 D,E). However, in osmotically stressed cells it lead to a ∼50% decrease in the surface accumulation of CI-M6PR (Fig. 8 D,E) in line with endocytosis and lysosomal exocytosis equally contributing to this effect. As a control, we tested the effect of the lysosomal exocytosis inhibitor on the surface level of the transferrin receptor little of which should reside at the lysosome. Indeed, we did not see any effect of lysosomal exocytosis inhibition on the surface level of transferrin receptor (Fig. 8F). In summary, a decrease in CI-M6PR endocytosis and strong surface accumulation upon osmotic stress, while its less pronounced enrichment at the plasma membrane upon heat stress is most likely exclusively due to the downregulation of its endocytosis.

### CI-M6PR contributes to cellular resilience upon osmotic stress

Our finding of the stress-induced accumulation of CI-M6PR at the plasma membrane raises the question whether this phenomenon is actually part of a stress response program rendering cells more resilient against osmotic stress or an inconsequential or even harmful side effect of stress-induced alterations in membrane transport. To address this question, we established an efficient knockdown of CI-M6PR in hTERT-RPE-1 cells (Fig. 9 A-D) and tested its effect on the survival and proliferation of stressed cells. As seen before, osmotic stress applied over several hours lead to an increase in the number of apoptotic and necrotic cells (Fig. 9 E,F). Depleting CI-M6PR similarly elevated the number of necrotic cells and tended to cause more apoptotic cells in line with its important cellular functions. However, more importantly, upon CI-M6PR depletion cells proved less resilient against osmotic stress with the number of necrotic and apoptotic cells roughly doubling in contrast to cells stressed while expressing CI-M6PR (Fig. 9 E,F).

**Fig. 9.**
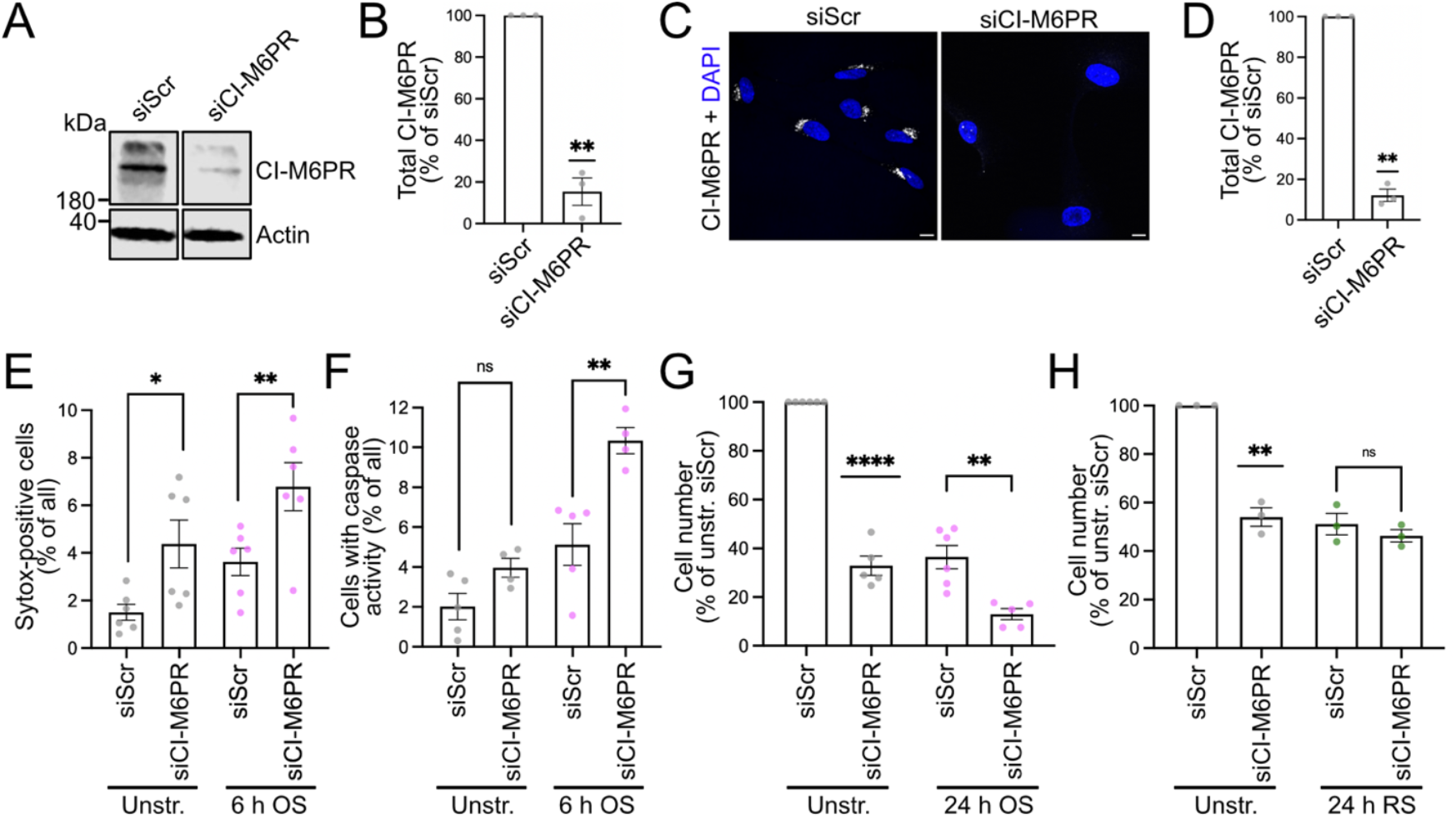
Cell death and proliferation decline induced by osmotic stress are further aggravated upon CI-M6PR silencing. *A-D*, efficient silencing of CI-M6PR by two rounds of siRNA transfection. *A*, representative Western blot. Bands originate from same blot. Aldolase A was used as loading control. *B*, quantification of CI-M6PR Western blot levels normalized to control (siScr). Analysis by one sample-test. *C*, representative confocal images of CI-M6PR knockdown cells stained for CI-M6PR. Nuclei were visualized by DAPI (in blue). Scale bars, 10 μm. *D*, quantification of the CI-M6PR immunofluorescence levels normalized to control (siScr). Analysis by one sample t-test. *E and F*, CI-M6PR silencing aggravates cell death upon osmotic stress. Silenced (siCI-M6PR) and control cells (siScr) were left unstressed or subjected for 6 h to 750 mOsm D-mannitol (OS) and subsequently incubated with SYTOX dye to label necrotic cells (E) or probed for caspase 3/7 activity to label apoptotic cells (F). Necrotic and apoptotic cells were quantified and expressed as % of all cells. Comparisons of control vs silenced cells under the different conditions by paired (E) resp. unpaired t-test (F) reveal a significant increase in stress-induced cell death upon CI-M6PR silencing. *G and H*, CI-M6PR silencing further decreases cell proliferation upon stress. Silenced and control cells were left unstressed or subjected for 24 h to 750 mOsm D-mannitol (OS; G) or 75 µM TBH (RS; H) and subsequently trypsinized and counted with the cell number expressed normalized to the unstressed control condition. Comparisons of unstressed control and silenced cells with each other by one-sample t-test show decrease in proliferation upon CI-M6PR silencing. Comparisons of stressed control and silenced cells by unpaired t-test demonstrate a significant additional decrease in cell proliferation in absence of CI-M6PR specifically upon osmotic stress. B, D-H: Columns depict mean ± SEM. Puncta indicate independent experiments. Ns, not significant; *=p<0.05, **=p<0.01, ****=p<0.0001. Details on n and p-values are provided in Source Data file.

When analyzing cellular proliferation upon stress, we again saw that hyperosmotic conditions reduced this process substantially, as did the downregulation of CI-M6PR (Fig. 9G). The negative effect of stress on proliferation was further exacerbated by CI-M6PR silencing. In fact, upon combined osmotic stress and CI-M6PR knockdown, the cell cultures attained only 10-15% of the cell number observed in unstressed non-silenced cells after 24 h of proliferation (Fig. 9G), indicating a pronounced downregulation of cell division. These findings strongly support the conclusion that CI-M6PR contributes to keeping cells fit to proliferate and that its loss is especially detrimental to cell proliferation under osmotic stress conditions. To validate the specificity of these findings for osmotic stress, the proliferation assay was repeated using oxidative stress, a condition which according to our surfaceome analyses does not induce CI-M6PR redistribution to the plasma membrane. Similarly to osmotic stress, oxidative stress reduced proliferation, however, there was no exacerbation upon CI-M6PR knockdown (Fig. 9H). This result supports our earlier conclusion that the detrimental impact of CI-M6PR depletion is particularly relevant upon osmotic stress when CI-M6PR is recruited to the plasma membrane.

### Testing CI-M6PRs role in lysosomal hydrolase sorting as cause for its protective effect

Since the best understood function of CI-M6PR is its role in escorting lysosomal hydrolases to late endosomes/lysosomes, we wondered whether its protective effect upon osmotic stress might be linked to this. The re-routing of CI-M6PR to the plasma membrane might potentially impair its ability to deliver hydrolases from the TGN to late endosomes. However, in light of the large intracellular pool of CI-M6PR, this scenario seems rather unlikely. On the other hand, increasing the residence time of CI-M6PR at the surface by downregulating its endocytosis, might promote the retrieval of lysosomal hydrolases, which appears especially relevant in light of the fact that we showed that osmotic stress leads to lysosomal exocytosis and thus to the acute loss of lysosomal hydrolases to the extracellular space.

As a first step to understand the impact of osmotic stress on the degradative capacity of hTERT-RPE-1 cells, we measured hydrolase activity with the dye Magic Red, which becomes fluorescent upon cleavage by cathepsin B. This revealed a decrease in intracellular hydrolase activity upon 1 h of osmotic stress (Fig. 10 A,B) consistent with the observed loss of hydrolases by lysosomal exocytosis (Fig. 8B), suggesting that the upregulation of CI-M6PR at the surface within this time frame cannot compensate for the loss of hydrolases. However, with ongoing osmotic stress the hydrolase activity levels returned to pre-stress levels (Fig. 10 A,B), maybe due to the surface-enriched CI-M6PR becoming effective in retrieving hydrolases and sorting them back to the endosomal system.

**Fig. 10.**
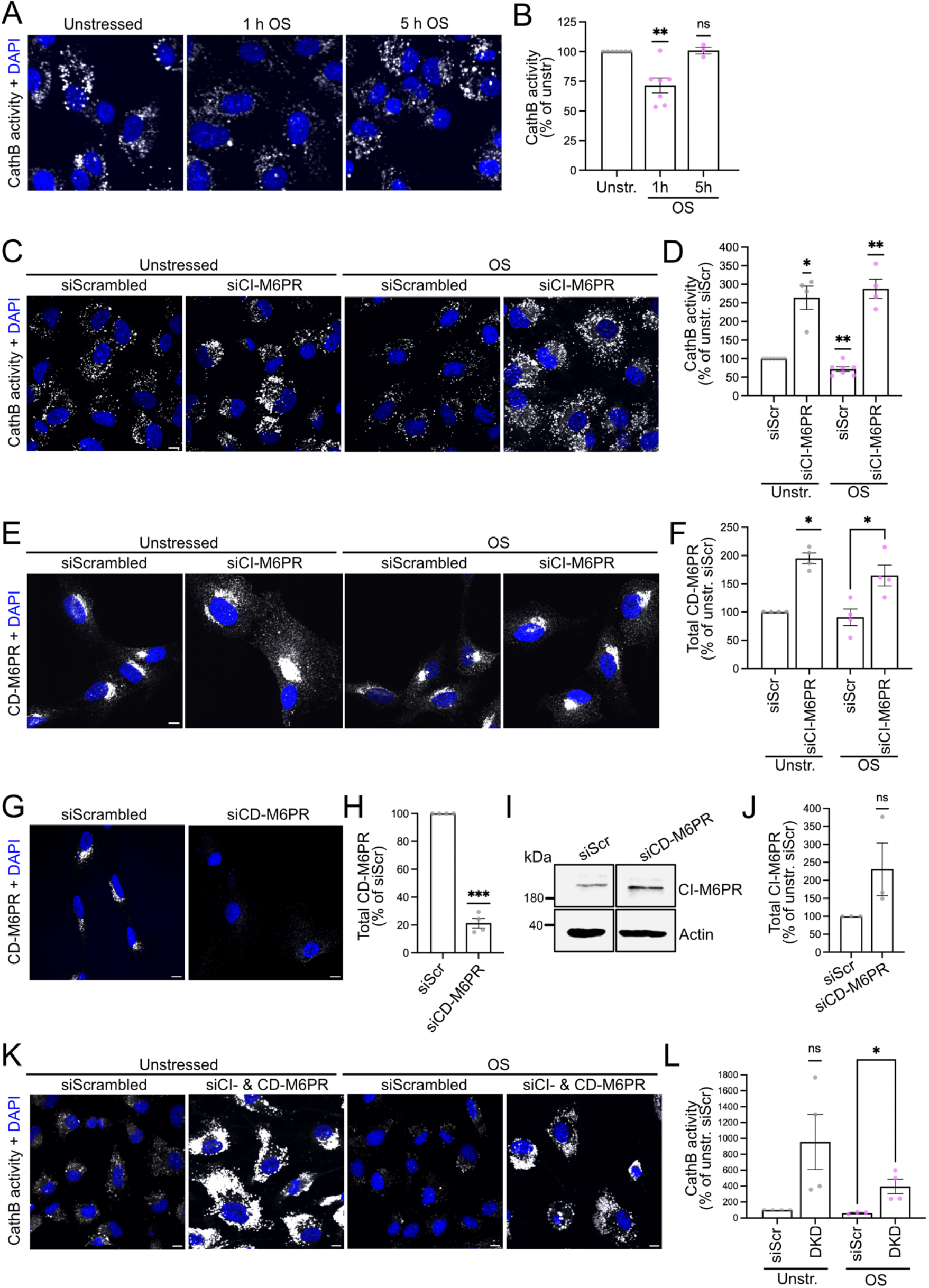
Stress-induced surface accumulation of CI-M6PR does not increase lysosomal degradative capacity. *A and B*, lysosomal hydrolase activity upon osmotic stress. Cells left unstressed or subjected to 750 mOsm D-mannitol (OS) for the indicated times were stained with Magic Red as indicator of Cathepsin B activity. *A*, representative confocal images. Nuclei were labelled with DAPI (in blue). Scale bar, 10 µm. *B*, quantification of Cathepsin B activity expressed as % of unstressed condition. Analysis by one sample t-test reveals a significant decline in hydrolase activity upon 1 h of osmotic stress which normalizes after 5 h. *C and D*, CI-M6PR silencing surprisingly increases hydrolase activity. Control (SiScr) and CI-M6PR knockdown cells (siCI-M6PR) were left unstressed or subjected for 1 h to 750 mOsm D-mannitol (OS) and subsequently stained with Magic Red. *C*, representative confocal images. Nuclei were labelled with DAPI (in blue). Scale bar, 10 µm. *D*, quantification of Cathepsin B activity expressed as % of unstressed control condition. Analysis by one sample t-test confirms decrease in hydrolase activity upon 1 h osmotic stress and reveals significant increase upon CI-M6PR silencing irrespective of osmotic stress. *E and F*, upregulation of CD-M6PR upon CI-M6PR silencing. Control and CI-M6PR knockdown cells were left unstressed or subjected for 1 h to 750 mOsm D-mannitol (OS) and subsequently stained with CD-M6PR-specific antibodies. Nuclei were labelled with DAPI (in blue). *E*, representative confocal images. Scale bar, 10 µm. *F*, quantification of CD-M6PR levels expressed as % of unstressed control condition. Comparisons of unstressed cells with each other by one sample t-test and of stressed cells with each other by unpaired t-test indicates a significant increase in CD-M6PR upon CI-M6PR silencing irrespective of osmotic stress. *G-J*, upregulation of CI-M6PR upon CD-M6PR silencing. *G and H*, efficient silencing of CD-M6PR by two rounds of siRNA transfection. *G*, representative confocal images of CD-M6PR knockdown cells stained for CD-M6PR. Nuclei were visualized by DAPI (in blue). Scale bars, 10 μm. *H*, quantification of the CI-M6PR immunofluorescence levels normalized to control (siScr). Analysis by one sample t-test. *I*, representative Western blot of CD-M6PR silenced cells probed for CI-M6PR and actin as loading control. Bands originate from same blot. *J*, quantification of CI-M6PR level normalized to actin level and expressed as % of unstressed control condition. Analysis by one sample t-test. *K and L*, combined CI- and CD-M6PR silencing still results in higher hydrolase activity. Control and CI/CD-M6PR double knockdown cells were left unstressed or subjected for 1 h to 750 mOsm D-mannitol (OS) and subsequently stained with Magic Red. *K*, representative confocal images. Nuclei were labelled with DAPI (in blue). Scale bar, 10 µm. *L*, quantification of Cathepsin B activity expressed as % of unstressed control condition. Comparisons of unstressed cells with each other by one sample t-test and of stressed cells with each other by unpaired t-test indicates in the latter case an increase in hydrolase activity upon CI/CD-M6PR double knockdown. B,D,F,H,J,L: Columns depict mean ± SEM. Puncta indicate independent experiments. Ns, not significant; *=p<0.05, **=p<0.01. Details on n and p-values are provided in in Source Data file.

To verify the overall importance of CI-M6PR for hydrolase sorting in hTERT-RPE-1 cells, we measured hydrolase activity upon CI-M6PR silencing. To our surprise, we observed a large increase in hydrolase activity in absence of CI-M6PR (Fig. 10 C,D) which also overrode the decrease in hydrolase activity seen upon 1 h of osmotic stress (Fig. 10 B). This is clearly at odds with the impairment of hydrolase delivery to lysosomes and the resulting decrease in lysosomal hydrolase activity that has been published for other CI-M6PR deficient cells (Takeda et al., 2019) and for cells in which CI-M6PR trafficking is disturbed (Arighi et al., 2004; Cui et al., 2019). However, CI-M6PR is not the only protein escorting hydrolases to the lysosome. The cation-dependent M6PR (CD-M6PR) also transports hydrolases from the TGN to lysosomes (Ghosh et al., 2003) and was previously suggested to be able to partially compensate for loss of CI-M6PR (Dittmer et al., 1998). Therefore, we tested potential compensatory effects of CD-M6PR in hTERT-RPE-1 cells. Indeed, the knockdown of CI-M6PR lead to an increase in the levels of CD-M6PR in unstressed as well as osmotically stressed cells (Fig. 10 E,F). Reversely, the efficient silencing of CD-M6PR (Fig. 10 G,H) induced an elevation in the protein level of CI-M6PR (Fig. 10 I,J) indicating a mutual compensation. However, even when we depleted both M6PRs, we still observed an increase in hydrolase activity that overrode the decrease triggered by 1 h of osmotic stress (Fig. 10 K,L). This is likely due to compensation by further proteins reported to be involved in hydrolase trafficking such as sortilin (Aguilera et al., 2023), which our proteomics data also indicates to be upregulated upon osmotic stress (Fig. 8A, gene name: SORT1). This suggests that an intact lysosomal degradative capacity is highly important to retinal pigment epithelium cells, whose physiological function is to phagocytose and degrade shed outer segments of photoreceptor cells (Young and Bok, 1969). In line with this, the functionality of the hTERT-RPE-1 lysosomal system appears to be maintained by compensatory upregulations of redundantly acting hydrolase transport proteins upon silencing of key factors for hydrolase transport. Possibly, the decreased endocytosis of CI-M6PR from the plasma membrane upon osmotic stress is another means of hTERT-RPE-1 cells to counteract the loss of hydrolases from the lysosome, which is caused in this scenario due to the stress-triggered lysosomal exocytosis.

## DISCUSSION

Stress-induced adaptations of membrane transport pathways and the resultant surfaceome changes remain an underexplored area. Our systematic approach revealed that already comparably mild osmotic, oxidative or heat stress leads to significant decreases in endocytosis (Fig. 3 B,D), while osmotic stress is unique in triggering in addition also lysosomal exocytosis (Fig. 8B). As our quantitative MS-based analysis of stress-induced surfaceome changes demonstrated, these acute membrane transport alterations substantially modify the plasma membrane composition (Fig. 4) - before transcription-dependent stress responses can even take effect.

In line with the distinct challenges that different stress conditions pose, the investigated stressors exert differential effects on the surface levels of individual transmembrane proteins (Fig. 4). This suggests that the applied stressors at the chosen dosage do not trigger a general impairment of endocytosis in line with the lack of changes in the abundance of endocytic components in our proteomic analysis (Supplementary Fig. 2), but rather regulate transmembrane proteins on an individual basis. While this apparently leads to a similar overall decrease in the amount of internalized proteins, it is largely different sets of transmembrane proteins that are affected by the different stressors.

In this context, our experiments uncovered that CI-M6PR is a highly stress-responsive protein which becomes partially redistributed to the plasma membrane upon heat stress and strikingly accumulates there upon osmotic stress (Fig. 4 A,B; Fig. 6 A-F). We showed that its strong surface accumulation upon osmotic stress is the combined effect of its lower endocytosis (Fig. 7E) and increased surface delivery by lysosomal exocytosis (Fig. 8E). These results lay the foundation for studying in how far the altered trafficking of CI-M6PR might be part of a stress response program which elevates the resilience of cells to cope with osmotic stress. Regarding the well understood function of CI-M6PR in the transport of lysosomal hydrolases, it is conceivable that its prolonged dwell time at the plasma membrane due to its decreased endocytosis rate may be beneficial for more efficiently retrieving lysosomal hydrolases lost from the cell due to stress-triggered lysosomal exocytosis. However, CI-M6PR also has a large number of additional extracellular ligands including IGF2 (Bohnsack et al., 2024; Morgan et al., 1987), heparanase (Wood and Hulett, 2008), granzyme B (Motyka et al., 2000), uPAR (Godár et al., 1999; Schiller et al., 2009), plasminogen, latent TGF*β* (Godár et al., 1999) and retinoic acid (Kang et al., 1997) for which the relevance of binding to CI-M6PR is often only partially understood (Gauthier et al., 2024). Therefore, it is also imaginable that the increased surface levels of CI-M6PR promote binding to one or multiple of these additional ligands and thereby strengthen signaling cascades with beneficial effects for cellular stress resilience. This will be an exciting area for further research and might also mean that even if the upregulation of CI-M6PR is a general cell-type independent feature, which will have to be tested, the cellular effects might still be cell-type specific depending on which ligands are present in the surrounding extracellular space.

In fact, our investigation of NHE7 highlights similarities across cell types, but also cell-type specific differences in membrane transport-based stress response programs. Consistent with our earlier data in primary murine astrocytes (Lopez-Hernandez et al., 2020b), we found NHE7 to be surface-enriched upon osmotic stress (Fig. 4A, Fig. 5 A,B). However, while we showed astrocytes to upregulate lysosomal hydrolase activity in an NHE7-dependent manner upon prolonged hyperosmotic stress (Lopez-Hernandez et al., 2020b), the degradative capacity of osmotically stressed hTERT-RPE-1 cells was maintained at the pre-stress level after a transient decrease (Fig. 10B), presumably in line with their high resilience against perturbances in hydrolase activity. Therefore, it will be important in the future to conduct similar studies in multiple cell types to be able to distinguish global stress response programs from cell-type specific adaptations.

Another important question raised by our research is how the stress-induced differences in membrane transport are brought about. It is very intriguing that lysosomal exocytosis is specifically induced upon osmotic stress. So far, lysosomal exocytosis is mostly known to be involved in plasma membrane repair and reported to be triggered by a local influx of Ca^2+^ at the damaged site which promotes lysosomal fusion with the plasma membrane (Gerasimenko et al., 2001; Rodriguez et al., 1997). At first glance, it is not clear why hyperosmotic stress, which leads to cell shrinkage and thereby decreases membrane tension, should induce a membrane repair response. However, lysosomal exocytosis has been recognized to fulfill other functions as well. One example is extracellular matrix remodeling (Jiang et al., 2026; Trojani et al., 2024), a role it actually shares with CI-M6PR, which can recruit heparanase (Wood and Hulett, 2008). However, further research is needed to determine whether lysosomal exocytosis is indeed part of an osmoprotective response or a side effect of osmotic stress-induced alterations. Mechanistically, it is easily conceivable that, similar to the wound repair scenario, also upon osmotic stress a plasma membrane proximal elevation in Ca^2+^ is the trigger for lysosomal exocytosis, since we have previously shown that the osmotic stress-triggered surface enrichment of NHE7 with its subsequent NHE7-mediated Na^+^ influx induces Ca^2+^ entry via the Na^+^/Ca^2+^ exchanger NCX1 (Lopez-Hernandez et al., 2020b).

For endocytosis, it is known that the uptake of specific cargoes can be modulated by rapid post-translational modifications of either endocytic adaptor or cargo protein. For example, phosphorylation triggers the *β*-arrestin-dependent endocytosis of GPCRs (Liu et al., 2025), while ubiquitination is the prerequisite for CALM-dependent internalization of GluA1 (Azarnia Tehran et al., 2022). In line with this, modifications preventing ubiquitination can inhibit endocytosis as shown for GluA1 (Azarnia Tehran et al., 2022). ARH is an example for an endocytic adaptor which needs to be post-translationally modified by S-nitrosylation to successfully engage with the endocytic machinery (Zhao et al., 2013), a modification that might well be responsive to the cellular redox state and thus to oxidative stress. While these principal mechanisms for regulating the internalization of individual transmembrane proteins are well understood, the signaling cascades connecting the initial stress sensors to the enzymes mediating the required post-translational modifications have largely remained elusive.

In the case of CI-M6PR it is known that its cytoplasmic tail harboring its sorting determinants (Ghosh et al., 2003) is phosphorylated by casein kinases on multiple serines (Rosorius et al., 1993). In addition, screens have uncovered 127 kinases and phosphatases which directly or indirectly affect CI-M6PR trafficking downstream of diverse signaling pathways (Adachi et al., 2009; Anitei et al., 2014), and Anitei et al. reported on eleven genes whose knockdown caused a surface accumulation of a GFP-CI-M6PR chimera (Anitei et al., 2014). The challenge will be to decipher which of these many regulators specifically affect the endocytosis of CI-M6PR and are themselves regulated by a stress-responsive signaling cascade.

In conclusion, it will be of great interest for the future to delve deeper into the stress-specific surfaceome changes we discovered since it will be essential to elucidate the specific signaling and membrane transport changes underlying the redistribution of individual transmembrane proteins to or from the plasma membrane. Finally, it will be even more crucial to decipher in each case the relevance of the altered protein trafficking in the context of stress response programs aimed at strengthening cellular resilience. By revealing the stress-responsive trafficking of CI-M6PR, a phenomenon that had previously gone unnoticed, our work opens a new line of inquiry into stress-dependent alterations in membrane transport with a strong potential for advancing our understanding of human disease mechanisms influenced by these stress conditions.

## MATERIALS AND METHODS

### siRNAs

The following siRNAs were used in this study:

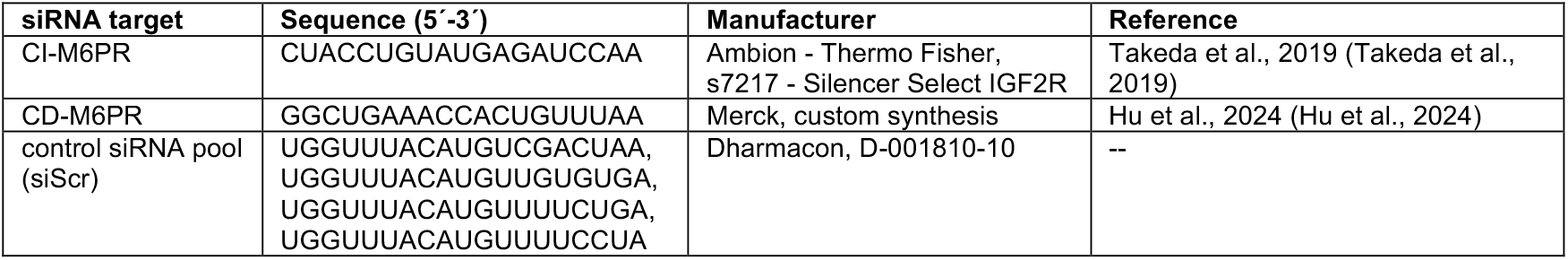

### Antibodies

The following primary antibodies were used in this study:

Antibodies The following primary antibodies were used in this study:
| Antigen | Host | Dilution Immunoblotting | Dilution Immunofluorescence | Manufacturer | Number |
| --- | --- | --- | --- | --- | --- |
| β-Actin | Mouse | 1:3000 | - | Sigma | A5441 |
| Aldolase A | Rabbit | 1:5000 | - | Proteintech | 11217-1-AP |
| α-Adaptin (AP-2α) | Mouse | 1:500 | 1:300 | BD Transduction | 610502 |
| CI-M6PR | Rabbit | 1:1000 | 1:400 | Cell Signalling | 14364S |
| CD-M6PR | Rabbit | - | 1:200 | Abcam | ab134153 |
| EEA1 | Mouse | 1:500 | 1:300 | Santa Cruz | sc-137130 |
| HSP70 | Mouse | 1:2000 | - | Thermo Fisher | MA3006 |
| SPPL2A | Rabbit | 1:1000 | 1:200 | Novus Biologicals | NBP2-93234 |
| TGN46 | Rabbit | - | 1:100 | Abcam | ab546 |
| Transferrin receptor | Rabbit | 1:500 | - | Sigma | HPA028598 |

The following secondary antibodies were used in this study (WB, Western blotting; IF, immunofluorescence):

| Antigen | Host | Conjugated with | Dilution | Manufacturer | Number |
| --- | --- | --- | --- | --- | --- |
| Mouse IgG | Goat | Alexa Fluor 488 | 1:200 (IF) | Thermo Fisher Scientific | A11029 |
| Rabbit IgG | Goat | Alexa Fluor 488 | 1:200 (IF) | Thermo Fisher Scientific | A11036 |
| Rabbit IgG | Goat | Alexa Fluor 647 | 1:200 (IF) | Thermo Fisher Scientific | A21244 |
| Mouse IgG | Goat | IRDye 680RD | 1:8000 (WB) | LI-COR Biosciences | 926-68070 |
| Mouse IgG | Goat | IRDye 800CW | 1:8000 (WB) | LI-COR Biosciences | 926-32210 |
| Rabbit IgG | Goat | IRDye 680RD | 1:8000 (WB) | LI-COR Biosciences | 926-68071 |
| Rabbit IgG | Goat | IRDye 800CW | 1:8000 (WB) | LI-COR Biosciences | 926-32211 |

### Probes and dyes

The following probes and dyes were used in this study according to the manufacturer’s instructions:

Probes and dyes The following probes and dyes were used in this study according to the manufacturer's instructions:
| Probe/dye name | Label for | Manufacturer | Number |
| --- | --- | --- | --- |
| Streptavidin, Alexa Fluor 647 | Biotinylated proteins | Thermo Fisher Scientific | S21374 |
| Streptavidin, Alexa Fluor 488 | Biotinylated proteins | Thermo Fisher Scientific | S11223 |
| Transferrin, Alexa Fluor 647 | Transferrin receptor | Invitrogen | T23366 |
| DAPI | Nucleus | Carl Roth | 6335.1 |
| Hoechst | Nucleus | Thermo Fisher Scientific | 62249 |
| PROTEOSTAT Aggresome detection kit | Protein aggregates | Enzo | ENZ-51035 |
| CellROX green | Oxidative stress | Thermo Fisher Scientific | C10444 |
| IncuCyte Cas 3/7 green | Apoptotic cells | Sartorius | 4440 |
| SYTOX Green | Necrotic cells | Thermo Fisher Scientific | S7020 |
| Magic Red Assay kit | Cathepsin B activity | Abcam | ab270772 |

### Plasmids

Human NHE7-mCherry was a kind gift of Prof. Volker Haucke (FMP, Germany). The GFP-CI-M6PR chimera comprising a signal peptide followed by GFP and the CI-M6PR transmembrane and cytosolic domain in pEGFP.C1 was a kind gift of Prof. Peter J. Cullen (University of Bristol, UK).

### Cell culture

The human retinal pigment epithelium cell line hTERT-RPE-1 was obtained from the American Type Culture Collection (ATCC). hTERT-RPE-1 cells were cultured in DMEM/F-12 supplemented with 10% (v/v) FBS and 1% (v/v) penicillin/streptomycin (referred to as full medium) in a cell culture incubator at 37°C and 5% CO_2_. All cells were routinely tested for and devoid of mycoplasma contamination.

### Proliferation assay

Cells were counted and equal numbers plated in fresh full medium into the wells of a 12-well plate. After 24 h (day1) the respective stress treatments were started. 24 h later (day 2), cells in each well were detached with TrypLE and counted in a Neubauer chamber. If the proliferation assay was done on silenced cells, the siRNA transfection was performed 4 days before reseeding into the 12-well plate.

### siRNA transfection

20,000 hTERT-RPE-1 cells/well were seeded into a 12-well plate. For siRNA transfection, Lipofectamine RNAiMAX was diluted 1:400 in Opti-MEM before addition of siRNA. After vortexing and 15 min incubation at room temperature (RT) the mixture was diluted 1:5 with full medium (final siRNA concentration: 9-10 nM). For all experiments, 2 rounds of knockdown were performed: a reverse transfection on day 1 and a forward transfection on the late afternoon of day 2. On day 3 the cells reveived fresh full medium. On the morning of day 4, cells were re-seeded either onto glass coverslips or into IBIDI chambers, and the final experiments were performed on day 5. For Western Blot lysates, 300,000 cells were seeded into 6 cm dishes and transfected with reagents scaled up to match the larger surface area. Cells were lysed four or five days after the first reverse transfection depending on cell confluency.

### Plasmid transfection

One day prior to transfection, cells were seeded onto Matrigel-coated glass coverslips in 12-well plates at a density of 80,000 cells/well in 700 µl full medium to reach 70–90% confluency on the following day. Per well 4 µl Lipofectamine 2000 were diluted in 100 µl Opti-MEM and incubated for 5 min at RT. In parallel, 1.6 µg plasmid DNA were diluted in 100 µl Opti-MEM. Both solutions were then combined, vortexed and incubated at RT for 20 min. The resulting transfection mix was added dropwise to each well. Cells were returned to the incubator for 5–6 h. Afterwards, the transfection medium was removed and replaced with fresh full medium. At least 24 h post-transfection, cells were processed for confocal microscopy.

### Induction of stress conditions

To prepare the medium for inducing osmotic stress, first the osmolarity of the full medium was measured using an osmometer (normally about 285-310 mOsm/kg, in short mOsm throughout the paper). Then the required amount of D-mannitol to reach the desired osmolarity was added to the full medium. Once the D-mannitol had dissolved, the osmolarity of the medium was re-measured for confirmation, before the medium was heated to 37°C and applied to the cells. To prepare the medium for inducing oxidative stress, a stock solution of 0.5 M tert-butyl hydroperoxide (TBH) in distilled water was diluted in full medium to the desired concentration (usually 75 µM). After vortexing and pre-heating to 37°C, the TBH-containing medium was applied to the cells. To induce heat stress, cells were incubated at 42°C. The length of the individual treatments is specified in the figure legends. Cells were kept in the cell culture incubator during the treatment.

### Immunofluorescence

Cells on glass cover slips were washed 1x with PBS containing 0.5 mM CaCl_2_ and 1 mM MgCl_2_ (PBS^2+^), placed on ice and fixed with ice-cold 4% paraformaldehyde (PFA) in PBS for 10 min. Afterwards, the cells were washed 3x in PBS, and blocked and permeabilized in goat serum dilution buffer (GSDB; 15% (v/v) goat serum, 0.25% (v/v) saponin in PBS) for 30 min at RT. Afterwards, cells were incubated (upside down) for 1 h at RT on droplets containing primary antibodies in GSDB (dilution see primary antibody table). Then the coverslips were washed 3x with PBS prior to a 30 min incubation with secondary antibodies (see secondary antibody table). After 3x washes with PBS, coverslips were incubated with 1 µg/ml DAPI in PBS for 10 min and mounted with ImmuMount.

### Treatment of cells with endocytosis inhibitor Pitstop2

Cells were seeded onto glass coverslips 24 h prior to treatment. A stock solution of 30 mM Pitstop2 (Abcam) in DMSO was freshly diluted in serum-free medium to 30 µM. Cells were washed with Dulbecco’s phosphate-buffered saline (DPBS) before adding serum-free medium containing either 30 µM Pitstop2 or a matching volume of DMSO. Cells were treated for at least 15 min in the respective medium. Transferrin uptake assays were conducted afterwards in the same medium containing the inhibitor or vehicle. Following treatment, cells were acid-washed for 2 min at 4°C in (0.1 M glycine, 0.02 M HCl, pH 3.0), washed with PBS, and fixed with 4% PFA in H_2_O for 10 min at 4°C. After three PBS washes, cells were stained with 1 µg/ml DAPI for 10 min, washed 3x with PBS and mounted with ImmuMount for imaging. For Western Blot experiments involving Pitstop2, the compound was added for 1 h.

### Aggresome detection assay

Protein aggregate formation was assessed using the Proteostat Aggresome Detection Kit. Cells were seeded directly onto 8-well glass-bottom chamber slides and cultured for 24-28 h. Afterwards, cell were either left untreated, subjected to osmotic stress or treated with 10 µM of the proteasome inhibitor MG-132 for 18 h as positive control. Cells were fixed for 10 min with 4% PFA in PBS at RT, followed by 3x PBS washes. Permeabilization was performed on ice for 30 min using the 0.3% Triton X-100 and 0.5% EDTA-containing solution in the commercial assay buffer. Cells were then washed with PBS and incubated for 30 min at RT in the dark with a dual-staining reagent containing the Proteostat detection dye and Hoechst 33342. After a final PBS wash, cells were imaged at 60x magnification using a confocal microscope.

### CellROX green assay

CellROX green which becomes strongly fluorescent upon oxidation and subsequently binds to DNA was used to measure oxidative stress in live cells. 20,000 cells/well were seeded into glass-bottom IBIDI chambers. The next day the cells were subjected to oxidative stress for 1 h at 37°C. The 2.5 mM CellROX stock solution in DMSO was diluted to a final concentration of 5 μM in full medium and 0.5% (v/v) of Hoechst dye were added. Cells were incubated for 30 min at 37°C in this medium, then washed 3x with DPBS and imaged immediately at 37°C/5% CO_2_ and 60x magnification using a confocal microscope. CellROX green fluorescence was quantified within the area of the DAPI-labelled nuclei.

### SYTOX cell death assay and IncuCyte Cas 3/7 apoptosis assay

20,000 cells/well were seeded into glass-bottom IBIDI chambers. After at least 24 h, cells were subjected to the different stress conditions for the times indicated in the figure legends. For the SYTOX assay, the medium was then replaced with fresh full medium containing 100 nM SYTOX and 0.5% (v/v) Hoechst 33342 dye to stain the nuclei of dead or all cells, respectively. For the apoptosis assay, the medium was changed to fresh full medium containing 5 μM IncuCyte Caspase 3/7 dye and 0.5% v/v of Hoechst 33342 dye to stain cells with active caspase 3/7 and label all nuclei. After 30 min incubation at 37°C, live cells were imaged at 37°C/5% CO_2_ and 20x magnification using a confocal microscope.

### Cathepsin B activity assay

Cathepsin B activity was measured using the Magic Red Cathepsin B Assay Kit. For experiments involving siRNA-mediated gene silencing, cells were first transfected on glass coverslips in 12-well plates and cultured for five days. The day before the assay, transfected cells were re-seeded into 8-well glass-bottom chamber slides. Cells were then subjected to stress treatments as indicated in the figure legends, while control groups remained untreated. Afterwards, cells were washed once with DPBS and incubated for 30 min at 37°C in the dark with fresh full medium containing Magic Red reagent (1:25) and 0.5% (v/V) Hoechst 33342. After incubation, cells were washed twice with DPBS, placed into Opti-MEM and imaged immediately at 37°C/5% CO_2_ and 40x magnification using a confocal microscope.

### Antibody feeding assay

To assess CI-M6PR internalization upon stress, cells were first subjected for 1 h to the different stress treatments and then immediately placed on ice. After one wash with ice-cold PBS^2+^, cells were incubated with anti-CI-M6PR antibodies (diluted 1:400 in full medium) for 30 min on ice to label surface-localized CI-M6PR. Unbound primary antibodies were removed by washing twice with PBS^2+^. To allow for endocytosis, coverslips were transferred to a pre-warmed humidified chamber and incubated at 37°C for 15 min. Control samples underwent instead a 2-min acid wash using cold glycine buffer (0.1 M glycine, 0.02 M HCl, pH 3.0) to remove residual surface-bound antibodies. After two PBS^2+^ washes, cells were fixed with 4% PFA in PBS at 4°C for 10 min and permeabilized/blocked with GSDB buffer for 15 min at RT. To visualize internalized primary antibodies, cells were incubated with Alexa Fluor 647-conjugated secondary antibody (diluted in GSDB buffer) for 30 min at RT. After three PBS washes, nuclei were counterstained by adding 1 µg/ml DAPI in PBS to the cells for 10 min. After three more PBS washes, coverslips were mounted with ImmuMount and allowed to dry for at least 24 h before imaging on a confocal microscope at 60x magnification.

### Microscopy-based analysis of the bulk uptake of biotinylated surface proteins and transferrin

Cells were first subjected for 1 h to the different stress treatments and then immediately placed on ice and washed with ice-cold PBS^2+^ to inhibit endocytosis. To label cell surface proteins, cells were incubated on ice for 30 min with 0.5 mg/ml sulfo-NHS-SS biotin. Excess biotin was quenched by two 3 min incubations in ice-cold glycine (50 mM) solution, followed by a wash with ice-cold PBS^2+^. Afterwards, cells were incubated at 37°C in pre-warmed full medium containing fluorescently labelled transferrin (25 μg/ml) to allow for endocytosis. After 15 min, endocytosis was halted by transfering cells into ice-cold PBS. Residual biotin was cleaved off by two 10 min incubations in MESNa solution (50 mM Tris, 100 mM NaCl, 1 mM EDTA, 0.2% BSA, pH 8.6), and transferrin remaining on the plasma membrane was removed with a 2-min acid wash (0.1 M glycine, 0.02 M HCl, pH 3.0) at 4°C. Cells were then washed with PBS^2+^ and fixed in pre-cooled 4% PFA in H_2_O for 10 min. Following fixation, cells were blocked/permeabilized with GSDB for 15 min at RT before incubation with 0.1 μg/ml fluorescently-labelled streptavidin in GSDB to detect internalized biotinylated proteins. After two washes 1 µg/ml DAPI in PBS was added for 10 min to label nuclei. After three more washes with PBS cells were mounted with ImmuMount and stored in the dark at 4°C until confocal imaging at 60x magnification.

### Image acquistion and analysis

Confocal imaging was performed with a Nikon spinning disk confocal microscope equipped with a photometrics MXR10015 camera, a Nikon PerfectFocus autofocus system and an Okolab environment control chamber for live cell imaging at 37°C and 5% CO_2_. A 20x air (Plan-Apo, NA 0.75, WD 1.0 mm), 40x air (Plan-Apo, NA 0.95, WD 0.25-0.16 mm) or a 60x oil immersion (Plan-Apo, NA 1.40, WD 0.13 mm) objective was used. The setup was controlled by the imaging-Software NIS (Nikon). Images were acquired as z-stacks (z=3, 0.5 μm per slide) and analyzed using FIJI. Fluorescence intensities of the channels of interest were quantified following the application of a Gaussian Blur filter (radius = 2 pixels) to reduce image noise. An automatic global thresholding algorithm (typically Otsu, Li, or Minimum) was then applied to segment the channel. The total number of cells per image was determined by quantifying the DAPI-stained nuclei, and fluorescence intensities were normalized to controls and expressed on a per-cell basis. For certain quantifications and colocalization analyses, the “ComDet” plugin in FIJI was employed. This object-based colocalization tool enables analysis of multi-channel images by allowing independent adjustment of object size and intensity thresholds for each channel. ComDet provides a summary output including object counts, colocalization statistics, and references to specific regions of interest (ROIs), along with visual overlays of segmented objects for validation.

### Treatment of cells with lysosomal exocytosis inhibitor

Cells cultured in 10 cm dishes until 85-90% confluency were treated (in absence or presence of osmotic stress) for 1 h at 37°C with 10 µM of the TRPML1 Ca^2+^ channel inhibitor ML-SI3 (provided in DMSO) in full medium. Vehicle controls received an equivalent volume of DMSO. Following treatment, all samples were placed at 4°C and processed for surface biotinylation as described further below.

### Lysosomal exocytosis assay based on measuring the activity of released *β*-hexosaminidase

Cells were seeded at 1.2×10^5^ per well in 12-well plates and cultured overnight. The following day, cells were washed once with ice-cold DPBS and incubated either with 250 µl NaCl buffer (120 mM NaCl, 5 mM KCl, 2 mM CaCl_2_, 2 mM MgCl_2_, 10 mM glucose, 12 mM Hepes, pH 7.4) containing or lacking freshly added lysosomal exocytosis inhibitor ML-SI3 (10 µM) for 7 min at 37°C on a pre-heated shaking plate, or under the different stress conditions in 400 µl phenol red–free medium for 1 h at 37°C. After stimulation, plates were immediately transferred to ice. Supernatants were collected and clarified by 11,000×g centrifugation for 10 min at 4°C to remove debris. Cells were gently washed twice with ice-cold DPBS and lysed in 300 µl TNE buffer (20 mM Tris pH 7.4, 150 mM NaCl, 1 mM EDTA, 1% NP-40, 5% glycerol, supplemented with 3 µl/ml protease inhibitor cocktail and 10 µl/ml PMSF) for 45 min on ice with gentle shaking. Lysates were vortexed briefly and clarified by 11,000×g centrifugation for 10 min at 4°C. β-Hexosaminidase activity was measured using 4-nitrophenyl-N-acetyl-β-D-glucosaminide (pNAG, Sigma) as substrate. Reactions were performed in 96-well plates in duplicate. For supernatant samples, 50 µl of clarified supernatant were mixed with 50 µl of 2 mM pNAG dissolved in 25 mM citrate buffer (pH 4.5). For cell lysates, 8 µl of lysate were diluted with 52 µl of PBS, and 50 µl of 2 mM pNAG dissolved in 25 mM citrate buffer (pH 4.5) were added to reach a total reaction volume of 110 µl. Samples were mixed gently and incubated for 2 h at 37°C. Reactions were terminated by adding 150 µl carbonate buffer (0.05 M Na_2_CO_3_/NaHCO_3_, pH 10), resulting in the development of a yellow color. Absorbance was measured at 405 nm using a microplate reader. Background absorbance was subtracted using blank wells containing either NaCl buffer (for supernatant samples) or TNE buffer (for lysate samples). Absorbance values from blank wells were subtracted from corresponding sample readings. Duplicate measurements were averaged. To account for differing sample input volumes, absorbance values were normalized to the proportion of total sample used in the reaction (supernatants: 50 µl of 250 µl total; lysates: 8 µl of 400 µl total). β-Hexosaminidase release was expressed as the ratio of normalized enzyme activity in the supernatant to the sum of activities in supernatant and lysate. Data was further normalized to unstimulated controls and presented as relative enzyme release (% of control).

### Cell lysis and Western blotting

Cells were placed on ice, gently rinsed with cold PBS^2+^ and harvested by scraping into ice-cold PBS^2+^. Cell suspensions were transferred into 1.5 ml Eppendorf tubes and centrifuged at 300×g for 3 min at 4°C. The resulting pellets were resuspended in lysis buffer (20 mM HEPES pH 7.4, 100 mM KCl, 2 mM MgCl_2_, 1% (v/v) Triton X-100, 1 mM PMSF, 0.3% (v/v) mammalian protease inhibitor cocktail (Merck, P8340)), vortexed for 10 s, and incubated for 30 min at 4°C on a rotator. Lysates were then cleared by centrifugation at 17,000×g for 10 min at 4°C. Supernatants were transferred to fresh tubes, and protein concentrations were determined using the Bradford assay. Samples were then mixed 1:1 with 2× Laemmli buffer, denatured at 92°C for 10 min and analyzed by SDS-PAGE and Western blotting using nitrocellulose membranes. Following transfer, membranes were briefly rinsed in Tris-buffered saline (TBS) and then blocked for 1 h at RT on a shaker using Intercept (TBS) blocking buffer (LI COR Biotech LLC). Blocked membranes were incubated with primary antibodies diluted in antibody dilution buffer (Intercept (TBS) blocking buffer diluted 1:1 with TBS containing 0.1% (v/v) Tween-20 (TBS-T) overnight at 4°C. The next day, membranes were washed 3x 3 min with TBS-T before incubation with secondary antibodies in antibody dilution buffer for 45 min at RT, followed by 3x 10 min washes with TBS-T. Washed membranes were imaged using the Odyssey Fc imaging system (LI COR Biotech LLC). Band intensities were quantified using Image Studio Lite software (LI COR Biotech LLC) and normalized to appropriate loading controls.

### Cell surface biotinylation and isolation of biotinylated proteins for Western blotting

Cells were seeded into 10 cm dishes to reach 90% confluency at the time of the experiment. After applying stress or control treatments, cells were placed on ice to inhibit endocytosis. After two washes with ice-cold PBS^2+^, 6 ml 0.5 mg/ml sulfo-NHS-SS biotin solution were added per dish (except for the no-biotin controls), and cells were incubated at 4°C for 30 min to label surface proteins. Unbound biotin was quenched by adding cold glycine solution (50 mM glycine in PBS^2+^) for 10 min, followed by a PBS^2+^ wash. Cells were then scraped into cold 400 µl PBS per dish. Cell suspensions were centrifuged, and pellets were resuspended in 300 μl 1x RIPA lysis buffer (Merck, 20-188) containing mammalian protease inhibitor cocktail (Merck, P8340), vortexed briefly, and incubated on a rotator at 4°C for 30 min. Lysates were vortexed again and centrifuged at 13,000 x g for 10 min. Pellets were discarded, and protein concentrations of the supernatants were determined using a Bradford assay at RT. Aliquots of the lysates were retained for Western blot analysis of total protein input. 80-100 µl Neutravidin bead suspension per sample were pre-washed 3x with RIPA buffer (without inhibitors) and then incubated overnight at 4°C on a rotator with lysates, adjusted to contain equal protein amounts (at least 550 μg) across experimental conditions. Using snap-cap spin columns to minimize bead loss, beads were washed for 5 min per wash with 1 ml of the following solution: 3 washes in RIPA buffer/1 M NaCl (1:1 vol/vol), and 3 washes in 2 M urea in 50 mM ammonium bicarbonate for western blotting experiments. For MS the number of washes was tripled. For elution for subsequent Western blotting, 55 μl pre-heated 2× Laemmli sample buffer with fresh 5% ß-mercaptoethanol were added to the beads. After 15 min at 95°C, beads were spun down, and the eluates were collected.

### Sample preparation of biotinylated protein for MS-based surfaceome analysis

Neutravidin bead-bound proteins were processed for MS using an on-bead digestion protocol. Beads were incubated in 50 μl 6 M urea-containing buffer (6 M Urea, 50 mM Tris-HCl pH 7.5, 5 mM DTT) and heated at 55°C to reduce disulfide bonds, followed by alkylation with 15 mM 2-chloroacetamide at RT in the dark with gentle shaking. To quench excess alkylating agent, samples were incubated in 5 mM DTT. The samples were then diluted with 50 mM Tris-HCl buffer to reduce the urea concentration below 2 M. 5 ng/μl Trypsin were added to the beads, and digestion proceeded overnight at 37°C in a humidified chamber. Peptides were recovered by centrifugation. After transferring the supernatants into new tubes, a second elution step with 50 μl 1.5 M urea-containing buffer was performed to extract residual peptides. Both eluates were combined, and digestion was stopped by acidifying with ∼25 μl 1% trifluoroacetic acid (TFA) to pH<2 (verified using pH indicator paper). Samples were centrifuged at 13,000xg for 10 min to pellet any insoluble material, and the supernatants were retained for desalting. For desalting, C18 stage tips were assembled using triple-layer C18 material packed into standard 200 µl pipette tips. Tips were activated by loading and later centrifuging off 100 μl of methanol and equilibrated by adding and later centrifuging off 100 μl of 0.1% aqueous formic acid (buffer A). Acidified peptide samples were then loaded onto the tips, and washed 2x with buffer A as previously described, and finally loaded with 60 μl buffer B containing formic acid and acetonitrile (0.1% formic acid, 80% acetonitrile). Elution was performed by centrifugation at 500xg. Eluates were dried in a SpeedVac concentrator at RT and subsequently resuspended in 12 μl of buffer A. Samples were sonicated briefly, centrifuged to remove particulate materials, and either stored at –20°C or analyzed immediately. Approximately 2 µl of each sample were analyzed by liquid chromatography (LC)-MS/MS using electrospray ionization on a Q-Exactive HF mass spectrometer. For total proteome analysis, a slightly different protocol was used, as detailed below.

### Sample preparation for MS-based whole-cell proteome analysis

Unstressed cells and cells having been subjected to the different stress treatments for 1 h were scraped into ice-cold PBS. The cell suspension was centrifuged at 3000 g for 3 min, and the supernatant was discarded. The cell pellets (∼10^6^ cells) were resuspended in 150 µl lysis buffer (6 M GdmCl, 10 mM TCEP, 40 mM CAA, 100 mM Tris pH 8.5 in MS grade water) and lysed by heating at 99°C and shaking at 1400 rpm for 10 min, followed by strong sonication (Branson sonifier: 1 min, 20% duty cycle, output 3) to shear DNA. Lysates were clarified by centrifugation at 10,000×g for 10 min, and supernatants were retained. Protein concentrations were determined using a BCA assay in a 96-well format by mixing 5 µl of sample or BSA standard in lysis buffer with water to a final volume of 15 µl, followed by the addition of 200 µL of BCA working reagent (50:1 solution A:B), an incubation at 37°C for 30 min, and spectrophotometric analysis. For digestion, 25 µg protein from each sample were adjusted to equal volume with lysis buffer, diluted 1:10 in digestion buffer (10% acetonitrile, 25 mM Tris pH 8.5 in MS grade water), and incubated overnight at 37°C with LysC and trypsin (1:50 each; 1 µl of 0.5 µg/µL trypsin per sample). Digests were acidified with TFA to a final concentration of 1% (pH <2) and centrifuged for 3 min at 12,000×g. ∼75% of the supernatant were loaded onto pre-equilibrated SDB-RPS stage tips (3-layer, 3M 2241). Stage tips were activated with 100 µl acetonitrile, equilibrated first with 100 µl 30% MeOH + 1% TFA, and then 0.2% TFA, followed by peptide loading, washing with 0.2% TFA, and elution with 60 µl of 80% acetonitrile + 5% ammonia. Eluates were dried for ∼40 min by speed-vac at RT, resuspended in 12 µl Buffer A^++^ (0.1 % formic acid, 0.01 % trifluoracetic acid), sonicated for 5 min in a water bath, briefly centrifuged, and either stored at −20°C or analyzed directly via LC-MS/MS (2 µl injection).

### LC-MS/MS analysis

Peptides (4 μl) were separated on 50 cm columns with an inner diameter of 75 µm packed in-house with C18 beads (Reprosil, 1.9 μm, Dr. Maisch) using an EASY-nLC 1200 system and eluting peptides were sprayed directly into a Q Exactive HF mass spectrometer in data-dependent acquisition mode. A 1.5 h or a 3 h linear gradient from 2% to 95% buffer B was applied for streptavidin pull-down or total proteome samples, respectively.

The surfaceome samples for osmotic and heat stress as well as controls were analyzed together (project P0199). The surfaceome samples for oxidative stress as well as controls were run together with samples of heat-stressed cells having a 4-hour recovery period (not analyzed in this paper) (project P0207).

All whole-cell proteome samples were analyzed together (project P0260; the fourth replicate of the heat stress condition failed quality control and was therefore excluded from all downstream analysis). Raw data were processed using MaxQuant (v2.6.4.0).

### Statistical analysis of MS Data

All MS data was analyzed in Perseus (Version 1.6.15.0) and results were visualized in R (Version 4.3.1). Log_2_ transformed label-free quantification (LFQ) intensities were filtered to exclude all protein groups annotated as “only identified by site”, reverse hits and potential contaminants. Surface proteins were annotated using the the Governa SURFME classifier for bona fide surface-resident membrane proteins (Governa et al., 2022). To identify significantly up- or downregulated proteins for each stress condition, protein groups were filtered to retain at least 3 valid values in either the stress or control group. Missing values were imputed using random values chosen from a normal distribution centered around the detection limit of the instrument (width 0.3; down shift 1.8; mode separate from each column). Two sample Student’s t-tests were performed, and p-values were adjusted for multiple testing using the Benjamini-Hochberg procedure. Significantly up- or downregulated proteins with a minimal fold change > 2 and an adjusted p-value < 0.05 were visualized as volcano plots and Venn diagrams using ggplot2 and VennDiagram. For principal component analysis and non-supervised hierarchical clustering, log_2_ LFQ intensities were filtered to retain 3 valid values in at least one replicate group. Missing values were imputed as described above and principal components were calculated with the R function prcomp(). Heatmaps were drawn using the function pheatmap().

### Statistics

All graphs depict mean ± SEM, and puncta represent independent experiments. Statistical analyses were performed with GraphPad Prism 10.6.1. Data was assessed for normal Gaussian distribution with the Shapiro Wilk normality test. Outlier analysis was performed based on the ROUT method. For comparisons of two normally distributed groups, either paired or unpaired two-sided t-tests were used. For comparisons of more than two groups of normally distributed data the one-way ANOVA was used with Dunnett’s multiple comparison test for comparing samples to a control group. For more than two groups of not normally distributed data, the Kruskal-Wallis test followed by the Dunn’s multiple comparison test was used. Data to be compared to a normalized control were evaluated using a one sample t-test when normally distributed or a Wilcoxon signed rank test when not normally distributed. Significance levels are indicated in the figures as *=p<0.05, **=p<0.01, ***=p<0.005, ****=p<0.0001, and ns=not significant. For raw data and details on p-values please see the Source Data file.

## Supporting information

Unprocessed Blots File

## Accompanying supplemental material

- The **Supplementary Figure 1** (at the end of this document) shows the principal component analysis and heat maps of the surfaceome data.
- The **Supplementary Figure 2** (at the end of this document) shows stress-induced changes in the whole cell proteome.
- The **Supplementary Figure “Unprocessed Blots”** contains uncropped versions of the immunoblots depicted in the figures.
- The **Source Data File** contains all data used for the statistical analyses presented in the figures.
- The **Supplementary Table 1** contains the processed surfaceome data for osmotic and heat stress plus unstressed and no-biotin controls.
- The **Supplementary Table 2** contains the processed surfaceome data for oxidative stress plus unstressed and no-biotin controls.
- The **Supplementary Table 3** contains the processed whole-cell proteome data for osmotic, heat and oxidative stress plus unstressed control.
- The **Supplementary Table 4** contains the data for the list of endocytic proteins depicted in Supplementary Figure 2.
- The **Supplementary Table 5** contains a list of differentially regulated proteins from the whole-cell proteome dataset which are shared between different stress conditions.

## Acknowledgements

We thank Ralph Reiss for technical assistance and the students Luwam Gebrezgi and Sophie Meinhold for performing select microscopical and biochemical experiments under the supervision of FM during their bachelor theses. This research was supported by grants of the Deutsche Forschungsgemeinschaft (DFG, German Research Foundation) to TM (RTG 2737 project A1 and project number 461336323) and by the Forschungsinitiative Rheinland-Pfalz BioComp to TM.

## Author contribution

FRM and TM conceived the project and designed the experiments. FRM performed all experiments except for MS measurements. ZS and MR contributed MS measurements and analyses. GG performed MS data analysis and prepared graphical data presentations. TM wrote the manuscript with input and approval from all authors.

## Author notes

The authors declare no competing interests.

---

**Fig. S1.**
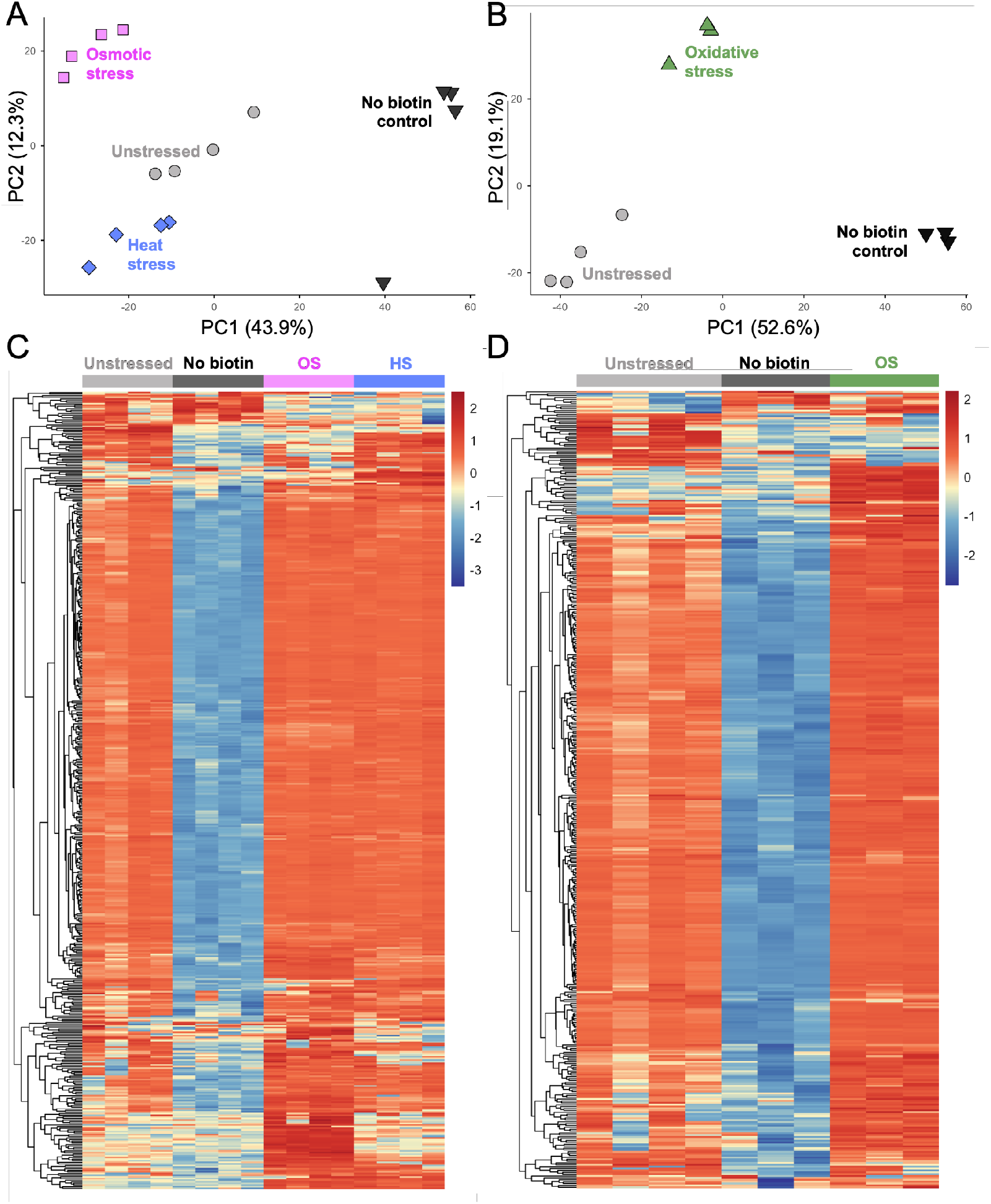
Principal component analysis and heat maps of surfaceome data. *A, B*, principal component analysis (PCA) of surfaceome data from the measurements of osmotic and heat stress samples plus unstressed control and no biotin control (A) as well as oxidative stress samples plus controls (B) showing separation between individual stress conditions and versus controls. Each symbol denotes an independently prepared sample. Samples analyzed within the same PCA were measured within the same MS experiment. *C, D*, Heatmap showing relative protein abundance across samples after Z-score normalization of surfaceome data from the measurements of osmotic and heat stress samples plus unstressed control and no biotin control (C) as well as oxidative stress samples plus controls (D) demonstrating consistency across replicates as well as successfull surface protein enrichment by biotinylation.

**Fig. S2.**
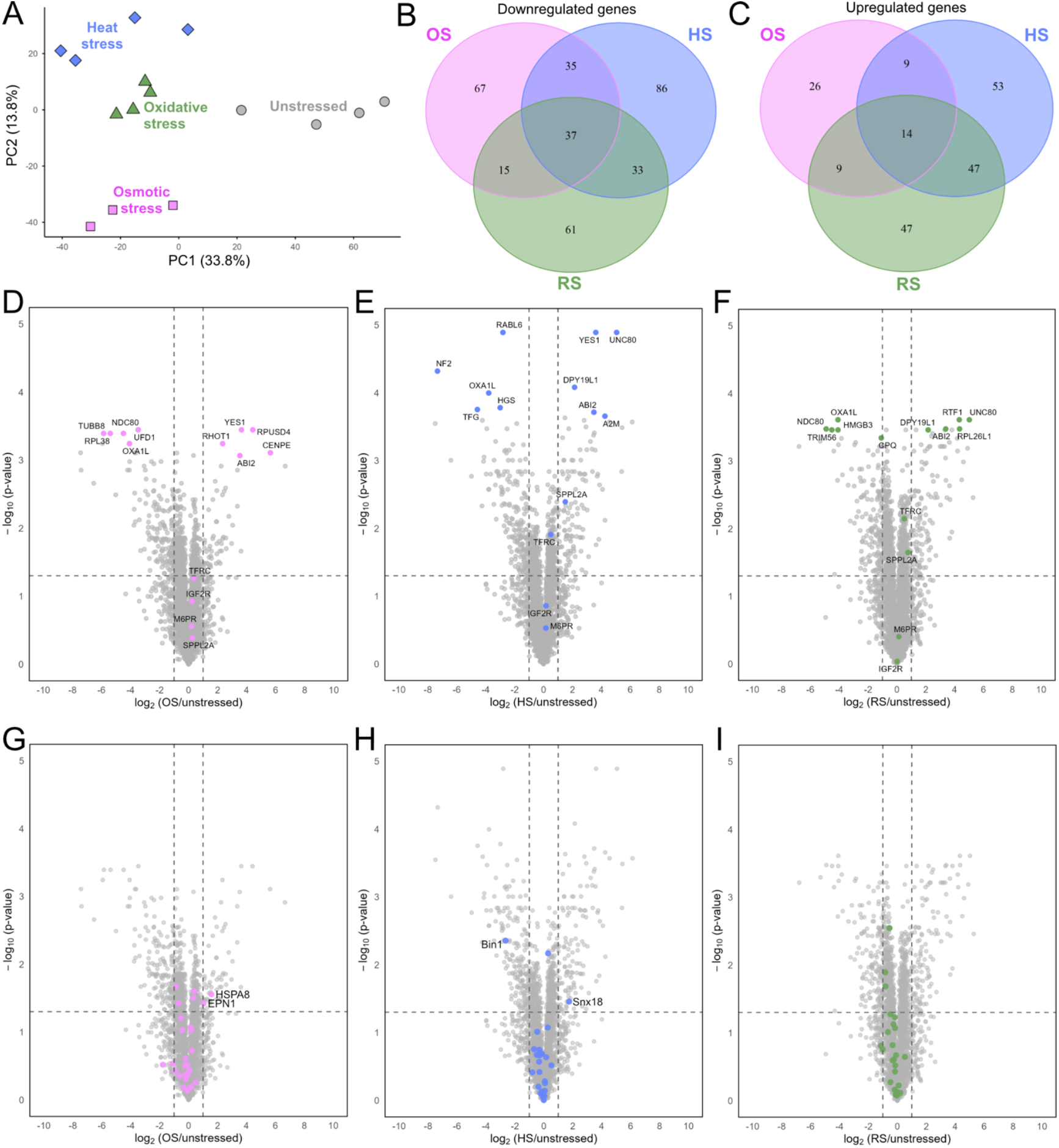
Stress-induced changes in the whole cell proteome. *A*, principal component analysis of whole-cell proteome data showing separation between individual stress conditions and versus control. Each symbol denotes an independently prepared sample. *B, C*, Venn diagrams visualizing the overlap in significantly deriched (B) or enriched (C) proteins across osmotic (OS), heat (HS), and oxidative stress (RS) conditions (BH-adjusted p<0.05, |log_2_FC|>1) (see also Supplementary Table 5). *D-F*, volcano plots of log_2_-fold changes and −log_10_-transformed, Benjamini–Hochberg corrected p-values representing whole-cell proteome alterations induced by OS (D), HS (E) or RS (F), highlighting the five most up- and the five most downregulated proteins according to p-value and log_2_-fold change as well as the validated candidates (see also Supplementary Table 3). *G-I*, volcano plots as above with endocytic proteins highlighted in colour (see also Supplementary Table 4). *D-I*, dashed lines represent the thresholds for statistically significant changes in protein abundance (threshold BH-adjusted p-value<0.05, |log_2_FC|>1).

