## Supplementary figures and images for "Analysis of stress-induced surfaceome remodeling reveals surface accumulation of the cation-independent mannose-6-phosphate receptor (CI-M6PR)"

### Unprocessed Blots File

Fig 1 E

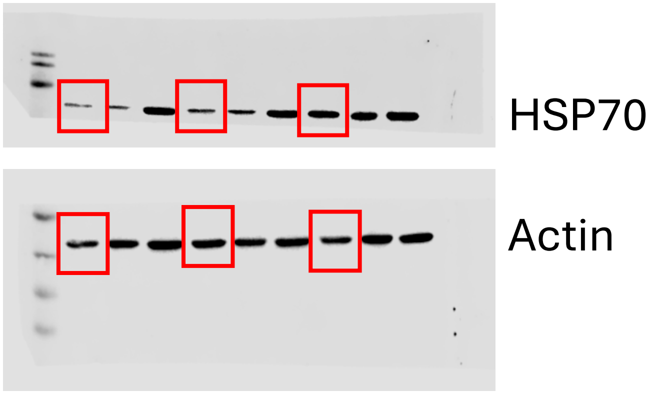


Fig 5 C

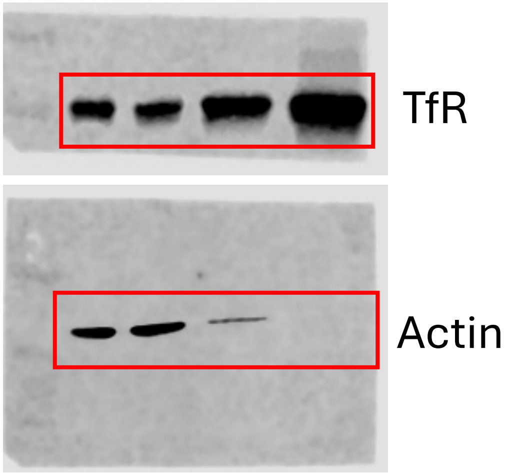


Fig 5 F

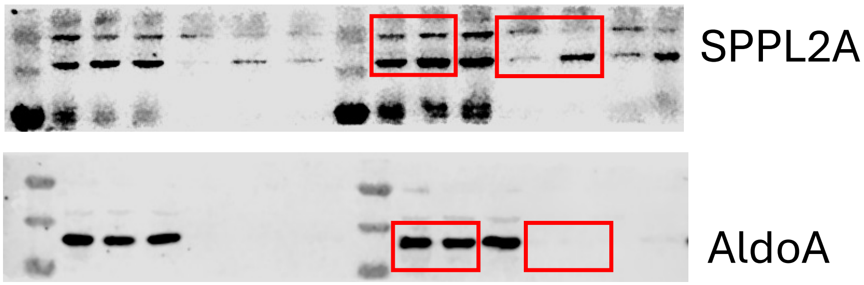


Fig 5 I

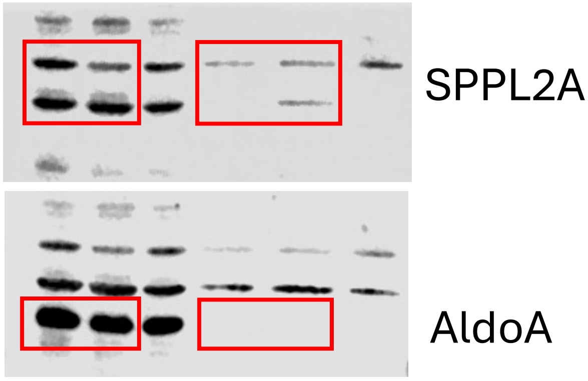

Fig 6 A

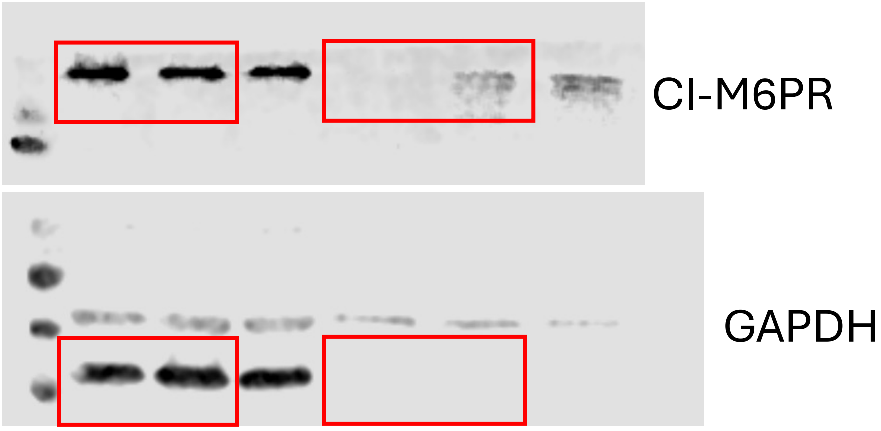


Fig 6 D

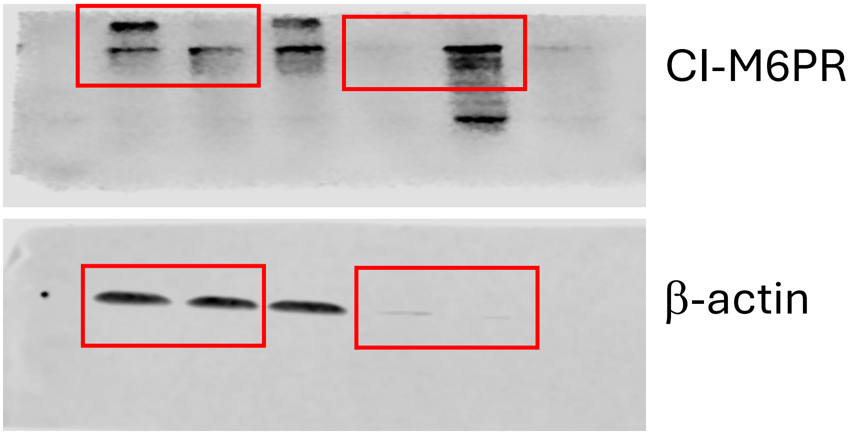


Fig 6 G

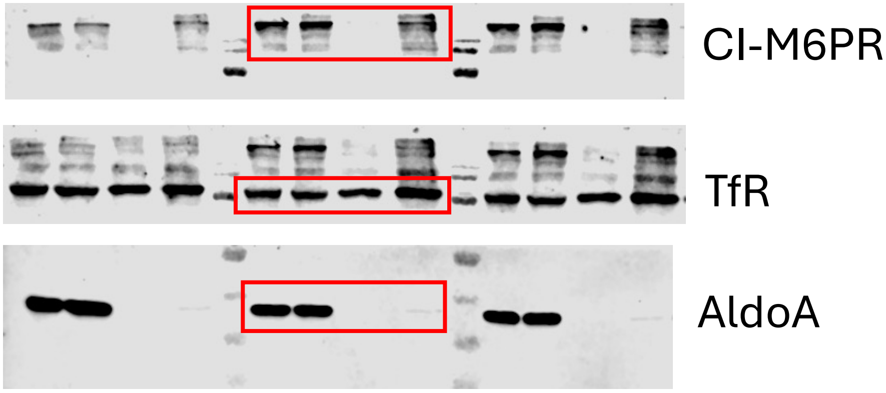

Fig 6 I

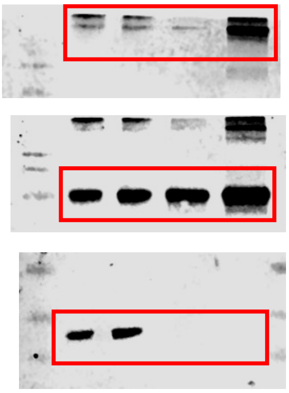


Fig 6 K

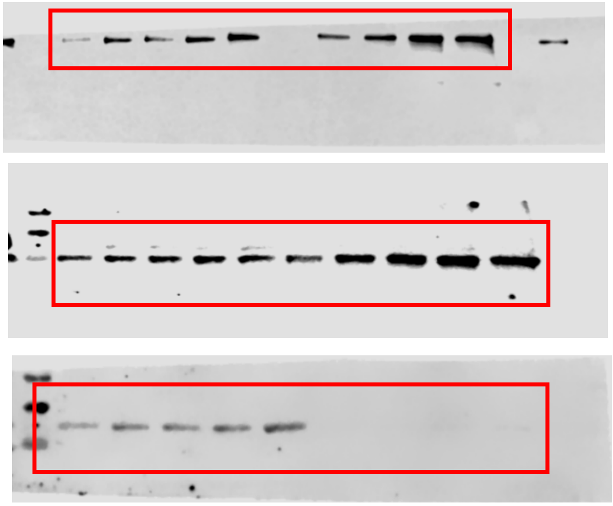


Fig 6 N

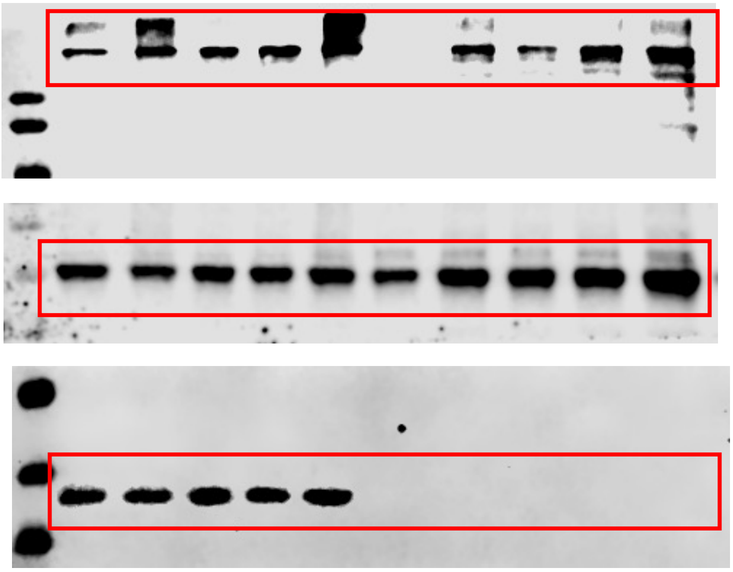

Fig 7 B

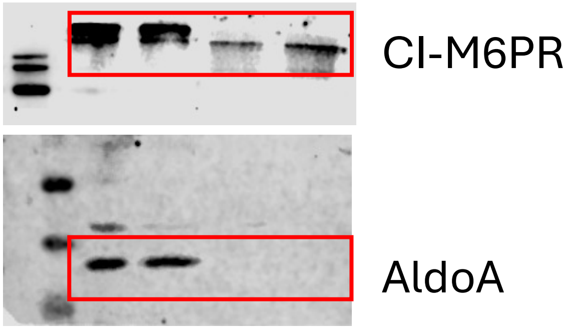


Fig 8 D

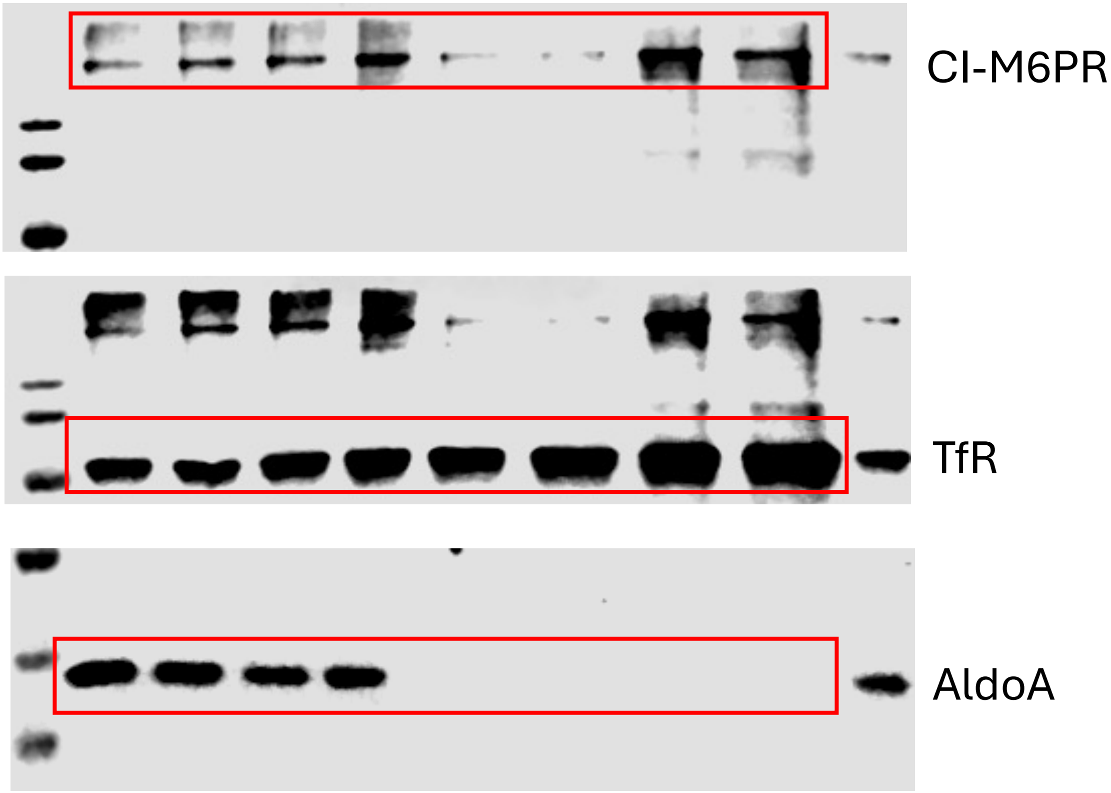


Fig 9 A

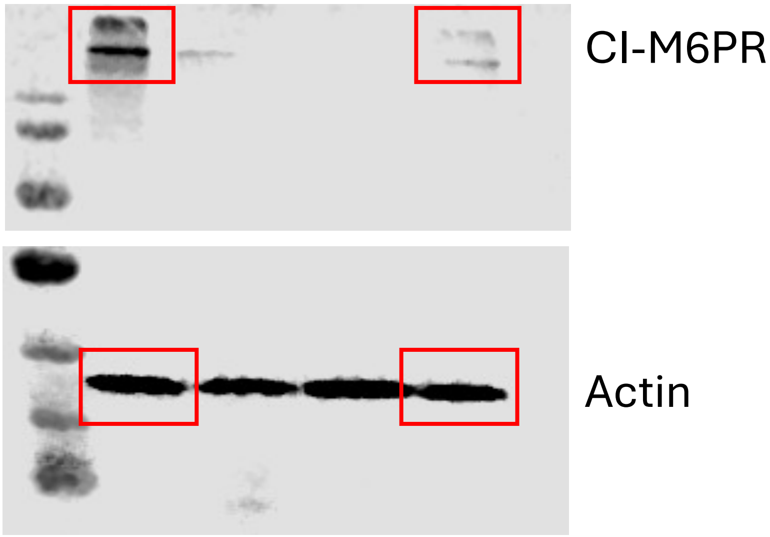

Fig 10 I

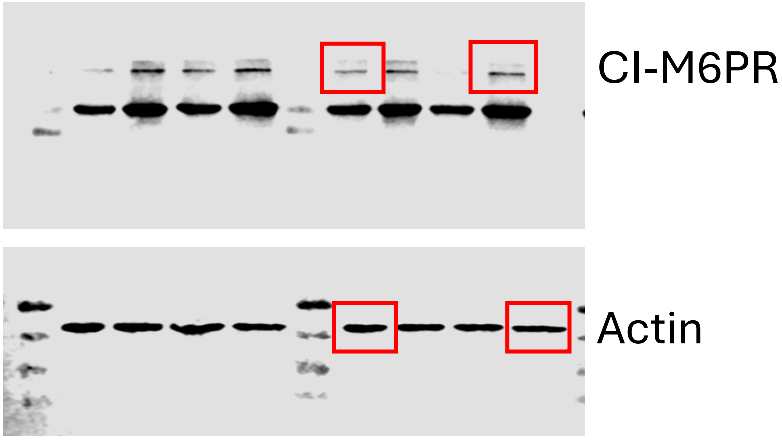
